# Cortical Betz cells analogue in songbirds utilizes Kv3.1 to generate ultranarrow spikes

**DOI:** 10.1101/2022.08.22.504741

**Authors:** Benjamin M. Zemel, Alexander A. Nevue, Leonardo E. Tavares, Andre Dagostin, Peter V. Lovell, Dezhe Z. Jin, Claudio V. Mello, Henrique von Gersdorff

## Abstract

Complex motor skills in vertebrates require specialized upper motor neurons with precise action potential (AP) firing. To examine how diverse populations of upper motor neurons subserve distinct functions and the specific repertoire of ion channels involved, we conducted a thorough study of the excitability of upper motor neurons controlling somatic motor function in the zebra finch. We found that robustus arcopallialis projection neurons (RAPNs), key command neurons for song production, exhibit ultranarrow spikes and higher firing rates compared to neurons controlling non-vocal somatic motor functions (AId neurons). This striking difference was primarily due to the expression of a high threshold, fast-activating voltage-gated K^+^ channel, Kv3.1 (KCNC1). RAPN properties thus mirror those of the sparse, specialized Betz cells in the motor cortex of humans and other primates, which also fire ultranarrow spikes enabled by Kv3.1 expression. These large layer 5 pyramidal neurons are involved in fine digit control and are notably absent in rodents. Our study thus provides evidence that songbird RAPNs and primate Betz cells have convergently evolved the use of Kv3.1 to ensure precise, rapid AP firing required for fast and complex motor skills.

## Introduction

The speed and accuracy of fine muscle control in mammals requires the firing of layer 5 pyramidal neurons (L5PNs) that project from the primary motor cortex (M1) to various targets in the brainstem and spinal cord (Oswald, Tantirigama, Sonntag, Hughes, & Empson, 2013; Suter, Migliore, & Shepherd, 2013). Studies in humans, macaques and cats have revealed a subclass of very large L5PNs called Betz cells, with wide-caliber myelinated axons, fast action potential (AP) conduction velocities and ultranarrow AP spikes (AP half-widths: 0.26 ms, *in vivo* recordings in macaques; 0.4 ms, brain slice recordings in cats, 35-37°C (Chen, Zhang, Hu, & Wu, 1996; Vigneswaran, Kraskov, & Lemon, 2011)). Betz cells were originally described in humans and they often terminate directly onto lower motor neurons (Lemon & Kraskov, 2019). They are thought to produce ultranarrow APs well suited for refined aspects of motor control. Remarkably, Betz cells are absent in rodent M1 cortex (Lacey et al., 2014; Soares et al., 2017). Studies of Betz-like cells have thus been hindered by species accessibility and because these specialized cells are interspersed with other smaller L5PNs in M1 cortex (Clustered in cotrical areas for fine digit control in humans; (Rivara, Sherwood, Bouras, & Hof, 2003; Tomasevic et al., 2022)).

Zebra finches are an accessible model organism for studying how upper motor neurons control a rapid and precise learned behavior. Their song consists of bouts of repeated motifs made up of multiple syllables, each containing distinct features that change on a rapid timescale (Leonardo & Fee, 2005; Sturdy, 1999; Yazaki-Sugiyama, Yanagihara, Fuller, & Lazarus, 2015). In sharp contrast with mammals, which possess a six layered neocortex, songbirds orchestrate song behavior with a circuitry consisting of pallial nuclei, which nonetheless have connectivity analogous to mammalian cortical microcircuits (Jarvis, 2019; Karten, 2015). Projection neurons within the robust nucleus of the arcopallium (RAPNs) represent the main forebrain output of the vocal motor system of songbirds. They share several molecular markers with mammalian L5PNs (Dugas-Ford, Rowell, & Ragsdale, 2012; Nevue, Lovell, Wirthlin, & Mello, 2020; Pfenning et al., 2014; Stacho et al., 2020). During song production, RAPNs exhibit robust burst-pause firing with high temporal precision (∼0.2 ms variance in the first spike latency; (Chi & Margoliash, 2001)). They are thus well poised to orchestrate the rapid movements of the syringeal muscles (∼4 ms to peak force of muscle twitch; (Adam & Elemans, 2020; Adam et al., 2021)).

We have recently shown that zebra finch RAPNs share many electrophysiological properties with mammalian L5PNs, including a hyperpolarization activated-current (I_h_), a persistent Na^+^ current and a large transient Na^+^ current (I_NaT_) with a short onset latency (Zemel et al., 2021). These properties facilitate the minimally adapting, regular spiking of APs (Almog, Barkai, Lampert, & Korngreen, 2018; McCormick, Connors, Lighthall, & Prince, 1985). A major difference, however, is that RAPNs exhibit an ultranarrow AP half-width (Zemel et al., 2021), a property also found in primate Betz cells (Lemon, Baker, & Kraskov, 2021). This unique feature of Betz cells is associated with high expression of the voltage gated potassium channel Kv3.1 subunit (Bakken et al., 2021; Ichinohe et al., 2004; Soares et al., 2017). The high-voltage, fast activation and fast deactivation of these channels have been shown to significantly narrow the AP waveform, thus facilitating high-frequency firing (Hong, Rollman, Feinstein, & Sanchez, 2016; Kaczmarek & Zhang, 2017; Rudy & McBain, 2001).

Similar to mammalian Betz cells, RAPNs in adult male finches also exhibit high expression of KCNC1 (Kv3.1; (Lovell, Carleton, & Mello, 2013)). Interestingly, this expression is significantly lower in the adjacent dorsal intermediate arcopallium (Aid) (Nevue et al., 2020), an area thought to be involved in non-vocal somatic motor functions (Feenders et al., 2008; Mandelblat-Cerf, Las, Denisenko, & Fee, 2014; Yuan & Bottjer, 2020) that have different temporal requirements than song (e.g., ∼10 ms to peak force of twitch for pectoral wing muscles (Bahlman, Baliga, & Altshuler, 2020)). Importantly, whereas the excitable properties of RAPNs have been examined in detail (Adret & Margoliash, 2002; Garst-Orozco, Babadi, & Olveczky, 2014; Liao, Hou, Liu, Long, & Li, 2011; Spiro, Dalva, & Mooney, 1999; Zemel et al., 2021), little is known about 1) the excitability of AId neurons, and 2) the molecular basis shaping the AP waveform in either brain region. Thus, a comparison of excitable features between RAPNs and AId neurons may reveal molecular specializations that evolved to specifically support the execution of complex, learned vocalizations.

Here we show that compared to RAPNs, AId neurons have broader APs that spike at lower frequencies during current injections. We found that these differences are due, in part, to the differential activity of Shaw-related K^+^ channels (Kv3.1-3.4), as Kv3 channel blockers disproportionately broadened APs and decreased firing rates in RAPNs compared to AId neurons.

Furthermore, a novel Kv3.1/3.2 positive modulator, AUT5, narrowed the AP waveform and increased the steady state firing frequency of RAPNs, but not of AId neurons. Moreover, KCNC1 had significantly higher expression in RA compared to AId, while KCNC2-4 (Kv3.2-3.4) genes were non-differential in expression. Notably, our analysis identified zebra finch KCNC3 (Kv3.3), a gene previously thought to be absent in birds. Morphological analysis revealed higher dendritic complexity and spine density in AId neurons compared to RAPNs, which also had larger spines, like Betz cells (Kaiserman-Abramof & Peters, 1972). We propose that the shared molecular and electrophysiological properties that promote ultranarrow spikes in songbird RAPNs and primate Betz cells likely originated from a convergent evolutionary process that allowed these neurons to operate as fast and precise signaling devices for fine motor control.

## Results

### Morphological features of RA and AId neurons

RA and AId are thought to control vocal and non-vocal somatic functions respectively via descending projections out of the finch telencephalon towards lower motor centers ((Bottjer, Brady, & Cribbs, 2000; Feenders et al., 2008; Mandelblat-Cerf et al., 2014; Nottebohm, Stokes, & Leonard, 1976; Paton, Manogue, & Nottebohm, 1981; Wild, 1993a; Yuan & Bottjer, 2020); Fig. 1A). Their close proximity in the arcopallium can be easily observed in frontal sections (Fig 1B-C; Fig. 2A-B). Whereas many morphological characteristics of RAPNs have been described, (Hayase et al., 2018; Kittelberger & Mooney, 1999; Miller, Cheung, & Brainard, 2017; Spiro et al., 1999), those of AId neurons have not been previously examined. Intracellular biocytin fills in frontal slices revealed that recorded AId neurons exhibit several morphological characteristics similar to those of RAPNs, including a large soma, a large diameter axon initial segment with extensively branched thin, aspinous axonal collaterals, and an extensive dendritic arborization, with individual dendrites exhibiting numerous spines (Fig. 1D-F). There were however some important differences. A Sholl analysis (Donald Arthur Sholl, 1956) revealed the extent of dendritic complexity for AId neurons to be significantly greater than that of RAPNs, although dendrites of AId neurons were restricted to ∼200 µm from the cell body, as has been previously reported for RAPNs (Fig. 1G). AId neurons also had approximately twice the number of spines compared to RAPNs as measured from a 30 µm segment of the tertiary branch of the filled cells (0.9 ± 0.07 vs. 0.4 ± 0.05 spines/µm; Fig. 1H). Additionally, whereas RAPN spines consistently presented with a mushroom-type shape, those in AId neurons exhibited a mix of mushroom, thin and filopodic structures (Fig. 1D-E, right; for a review on spine shapes see (Pchitskaya & Bezprozvanny, 2020).

**Figure 1.**
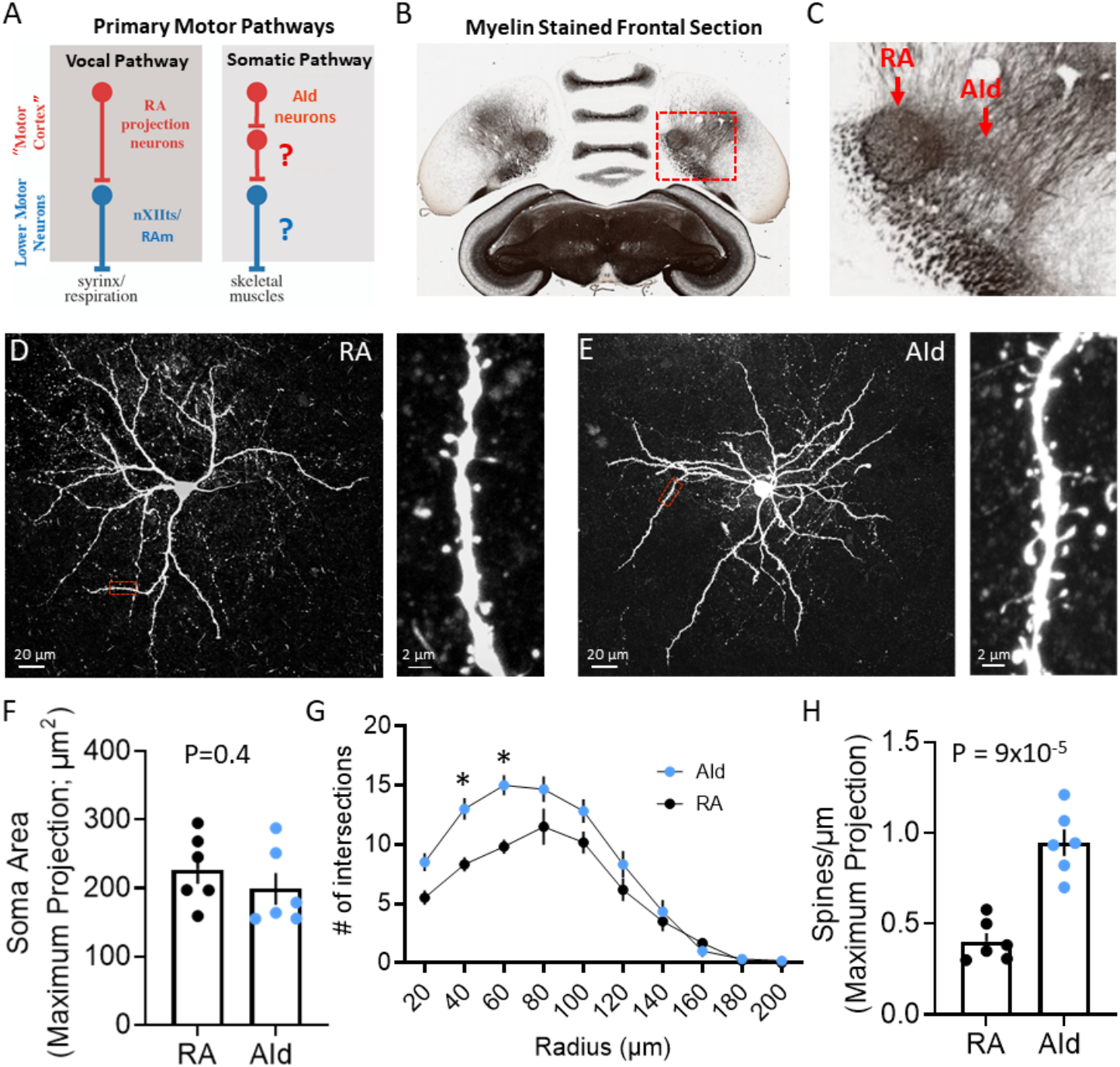
Morphology of RAPNs and AId neurons. A. Diagram of the motor pathways in the zebra finch. RA and AId upper-motor neurons in the arcopallium project to downstream targets that innervate syringeal and skeletal muscles respectively. Note that the exact targets for AId neurons in the brainstem (Bottjer et al., 2000) and/or spinal cord have yet to be determined. B. Myelin-stained frontal section of a zebra finch brain, including RA and AId (adapted from Karten et al. (Karten et al., 2013)). Red box in (B) indicates the region containing RA and AId. C. Expanded region shown in (B). Note the clear RA and AId arcopallial regions with inputs from the nidopallium above. D. Left: Maximum projection of a z-stack from a biocytin-filled RAPN. Red box indicates dendritic region expanded on the right. Right: Expanded region showing a 30 µm long dendritic section from the neuron to the left. E. Left: Maximum projection of a z-stack from a biocytin-filled AId neuron. Red box indicates dendritic region expanded on the right. Right: Expanded region showing a 30 µm long dendritic section from the neuron to the left. F. Dot plot comparing the soma area of RAPNs and AId neurons. Student’s t-test, two-tailed, t_stat_ = 0.91, N= 6 RAPNs and 6 AId neurons. Black bars indicate standard error. G. Comparison of the dendritic complexity from RAPNs and AId neurons as determined by the number of intersections with concentric circles of increasing radii surrounding the cell body. Two-way ANOVA; P = 0.004, F (9, 100) = 2.92, N = 6 RAPNs and 6 AId neurons. * P < 0.05 as determined by Tukey’s post-hoc test. Black bars indicate standard error. H. Dot plot comparing the number of spines per micron of RAPNs and AId neurons. Spines were counted on a 30 µm dendritic segment after the second branch point. Between 2-4 segments were measured per cell, averaged and divided by 30 to determine the # spines/µm. Student’s t-test, two-tailed, t_stat_ = 6.29, N= 6 RAPNs and 6 AId neurons. Black bars indicate standard error.

**Figure 2.**
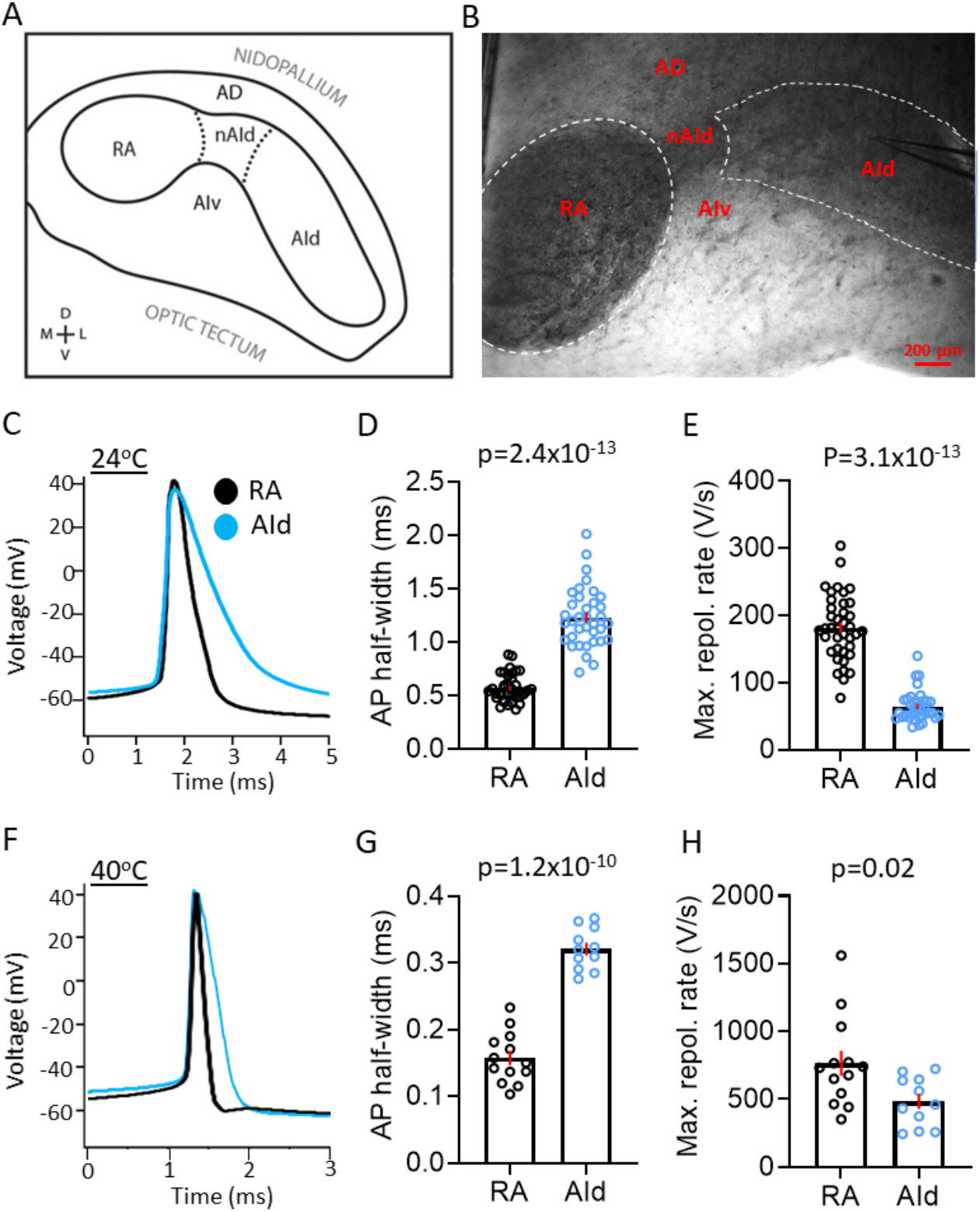
Spontaneous AP properties in RAPNs and AId neurons from adult male zebra finches at room and physiological temperatures. A. Drawing of the zebra finch arcopallium in the frontal plane, depicting RA and AId as defined by SCN3B staining in adult males (Nevue et al., 2020). Labeling of other domains are included as referential landmarks. *AD* dorsal arcopallium, *AId* dorsal intermediate arcopallium, *AIv* ventral intermediate arcopallium, *nAId* neck of the dorsal intermediate arcopallium, *RA* robust nucleus of the arcopallium. B. IR-DIC image of a frontal brain slice from which RA and AId are clearly visible. Labels correspond to those depicted in (A). Note the recording electrode in AId. C. Overlay of averaged spontaneous APs recorded at ∼24°C from a RAPN (black) and AId neurons (blue). D. Comparison of the average spontaneous AP half-widths from RAPNs and AId neurons recorded at ∼24°C (Mann-Whitney *U*= 11, two-tailed, N= 39 RAPNs and 36 AId neurons. Red bars indicate standard error). E. Comparison of the average spontaneous AP maximum repolarization rate from RAPNs and AId neurons recorded at ∼24°C. Mann-Whitney *U*= 14, two-tailed N= 39 RAPNs and 36 AId neurons. Red bars indicate standard error. F. Overlay of averaged spontaneous APs recorded at ∼40°C from a RAPN (black) and AId neurons (blue) respectively. G. Comparison of the average spontaneous AP half-widths from RAPNs and AId neurons recorded at ∼40°C. Student’s t-test, two-tailed, t_stat_ = 11.33, N= 13 RAPNs and 11 AId neurons. Red bars indicate standard error. H. Comparison of the average spontaneous AP maximum repolarization rate from RAPNs and AId neurons recorded at ∼40°C. Student’s t-test, two-tailed, t_stat_ = 2.50, N= 13 RAPNs and 11 AId neurons. Red bars indicate standard error. P-values are included in the graphs.

To gain further insights into the morphology of RAPNs and AId neurons we next analyzed our highest quality images (3 RAPNs and 2 AId neurons) using ShuTu software designed for digital 3-D reconstructions (Supplementary Fig. 1A-B; Video 1-2; (Jin et al., 2019)). We were able to confirm the approximately two-fold larger spine density when pooling all branch segments together (Supplementary Fig. 1C; Supplementary Table 1). The distribution of these spines as a function of distance from the soma was unimodal in both groups, with an increase in spines occurring after the initial, mostly aspinous primary dendrite, followed by a decrease toward more distal branches (Supplementary Fig. 1E). Upon manually tracing >1000 individual spines for each group we found that AId neurons display a larger proportion of spines with smaller 2-D surface areas compared to RAPNs (Supplementary Fig. 1D). The total surface area of the soma and dendrites of RAPNs and AId neurons is shown in Supplementary Table 1. The percentage of total surface for different compartments are soma: 13.4% vs. 9.5%; dendrites: 76.9% vs. 72.6%; spines: 9.7% and 17.9%, for RAPN and AId neurons, respectively (axons not included; Supplementary Table 1). While we acknowledge the limited data set and the caveat that some dendrites are cut off by the slicing procedure, the results suggest that AId neurons may receive more numerous, perhaps diverse, synaptic inputs that are differentially filtered compared to RAPNs. Finally, we calculated the surface to volume (S/V) ratio. For RAPNs we find 1.5-2.3 µm^-1^ and for AId neurons 2.5-2.9 µm^-1^ (Supplementary Table 1). These values are similar to those found using EM 3-D reconstructions in mouse cortical neurons (1.63 µm^-1^) but lower than found in astrocytes (4.4 µm^-1^ (Cali et al., 2019)), which may have metabolic energy consequences for RA nuclei, which stain heavily for metabolic markers (Adret & Margoliash, 2002).

### RAPNs and AId neurons: Passive properties

Fig. 2A depicts the male zebra finch arcopallium region in the frontal plane containing RA and AId brain regions, while Fig. 2B shows an infrared differential interference contrast (IR-DIC) microscopy image of a brain slice containing both brain regions. Note the heavily myelinated areas containing RA (Champoux, Miller, & Perkel, 2021) and AId. A comparison of RAPNs and AId neurons recorded in frontal sections at both room temperature (24°C) and physiologically relevant temperature (40°C (Aronov & Fee, 2012)) in the whole cell current clamp configuration revealed only minor differences in several passive membrane properties, including the membrane time constant, input resistance and the calculated membrane capacitance (C_m_) (Table 1). Using the calculated C_m_ from our room temperature recordings, we determined that the estimated average surface area was not significantly different between RAPNs and AId neurons (Mean ± SEM: 11563.6 ± 2277.5 µm^2^ vs. 7578.6 ± 1564.7 µm^2^ for RAPNs and AId neurons, respectively; Student’s t-test, t_stat_ = 1.331, P = 0.2). When considering possible tissue shrinkage (≤ 20%) during processing for fluorescent imaging (Winsor, 1994), these values approximate those found from results obtained from ShuTu total surface area reconstructions (Adjusted Mean: 8207.5 µm^2^ vs 9031.9 µm^2^ (including the soma, dendrites and spines) for RAPNs and AId neurons, respectively; Supplementary Table 1).

**Table 1.** Passive and spontaneously active properties of adult male RAPNs and AId neurons. Statistical analysis comparing each temperature respectively. Statistics were performed using an unpaired, two-tailed, Student’s t-tests if data was normally distributed with equal variances. Otherwise a Mann-Whitney U test was used. Data shown as Mean ± SEM. Room temperature comparisons: AP #-P = 0.7, t_stat_ = 0.3825, AP thresh-P = 0.09, t_stat_ = 1.696, AP amp- P = 0.05, t_stat_ = 1.971, AP halfwidth- P = 2.4×10^−13^, U = 11, Max. depol. rate- P = 4.0×10^−6^, t_stat_ = 4.988, Max. repol. rate- P = 3.1×10^−13^, U = 14, AP peak- P = 0.2, t_stat_ = 1.255, AHP- P = 0.0001, t_stat_ = 4.090, Time constant- P = 0.5, t_stat_ = 0.7259, Mem. Capacitan- P = 0.2, t_stat_ = 1.326, Input resistance- P= 0.3, t_stat_ = 1.160. High temperature comparisons: AP #- P = 0.0002, U = 14, AP thresh- P = 0.7, t_stat_ = 0.3301, AP amp- P = 0.2, t_stat_ = 1.302, AP halfwidth- P = 1.2×10^−10^, t_stat_ = 11.33, Max. depol. rate- P = 0.6, t_stat_ = 0.4644, Max. repol. rate- P = 0.02, t_stat_ = 2.504, AP peak- P = 0.1, t_stat_ = 1.568, AHP- P = 0.006, t_stat_ = 3.025, Time constant- P = 0.01, t_stat_ =2.735, Mem. Capacitan- P = 0.3, t_stat_ = 1.027, Input resistance- P = 0.004, t_stat_ = 3.280. * = P < 0.05. Abbreviations- AP #: AP spikes produced per second, AP thresh.: AP threshold, AP amp.: AP amplitude as measured from the peak of the after-hyperpolarization to the AP peak, Max. depol. rate: Maximum depolarization rate derived from the AP phase plane plot. Max. repol. rate: Maximum repolarization rate derived from the AP phase plane plot, AHP: Peak of the AP after- hyperpolarization, Mem. capacitan.: Membrane capacitance.

|  | RA (24°C) | AId (24°C) | RA (40°C) | AId (40°C) |
| --- | --- | --- | --- | --- |
| AP # | 13.6 ± 1.71<br>N=39 | 12.9 ± 1.14<br>N=36 | 41 ± 6.70<br>N=13 | 13.2 ± 2.47<br>N=11 |
| AP thresh. (mV) | -55.3 ± 0.66<br>N=39 | -53.7 ± 0.62<br>N=36 | -52.0 ± 1.00<br>N=13 | -52.7 ± 1.71<br>N=11 |
| AP amp. (mV) | 92.8 ± 1.53<br>N=39 | 89.0 ± 1.39<br>N=36 | 96.3 ± 3.57<br>N=13 | 102.8 ± 3.28<br>N=11 |
| AP halfwidth (ms) | 0.57 ± 0.02*<br>N=39 | 1.23 ± 0.05<br>N=36 | 0.16 ± 0.01*<br>N=13 | 0.32 ± 0.01<br>N=11 |
| Max. depol. rate (V/s) | 658.4 ± 26.53*<br>N=39 | 477.1 ± 24.59<br>N=36 | 1288.5 ± 67.11<br>N=13 | 1339.4 ± 85.34<br>N=11 |
| Max. repol. rate (V/s) | 181.9 ± 7.75*<br>N=39 | 64.5 ± 3.71<br>N=36 | 764.7 ± 92.41*<br>N=13 | 482.3 ± 52.83<br>N=11 |
| AP peak (mV) | 37.4 ± 0.96<br>N=39 | 35.5 ± 1.19<br>N=36 | 44.5 ± 2.87<br>N=13 | 50.3 ± 2.09<br>N=11 |
| AHP (mV) | -71.9 ± 0.53<br>N=39 | -68.4 ± 0.67<br>N=36 | -65.7 ± 0.92<br>N=13 | -69.2 ± 0.61<br>N=11 |
| Time constant (ms) | 17.4 ± 1.9<br>N=8 | 15.2 ± 2.3<br>N=6 | 17.2 ± 1.36*<br>N=10 | 13.3 ± 0.60<br>N=12 |
| Mem. capacitan. (pF) | 115.4 ± 22.77<br>N=8 | 75.8 ± 15.65<br>N=6 | 96.1 ± 7.39<br>N=10 | 106.8 ± 7.44<br>N=12 |
| Input resistance (MΩ) | 176.6 ± 22.11<br>N=8 | 209.2 ± 13.03<br>N=6 | 185.2 ± 15.02*<br>N=10 | 129.08 ± 8.81<br>N=12 |

### RAPNs fire ultranarrow spikes at higher frequencies than AId neurons

RAPNs produce remarkably narrow APs (half-width = ∼0.2 ms at 40°C; (Zemel et al., 2021)), making them well suited for orchestrating rapid movements of the syringeal and respiratory muscles required for song production. Upon examining properties of spontaneous APs, we found that RAPNs and AId neurons shared similar threshold, amplitude, peak and after-hyperpolarization at both temperatures examined (Fig. 2C; Table 1). However, AId neurons had an AP half-width that was twice as broad as the RAPN APs (Fig. 2C-D and 2F-G; Table 1). The shorter half-width of RAPNs could be due to either a faster AP depolarization and/or repolarization rate. To determine which phase of the AP was responsible for this difference, we derived phase plane plots from averaged spontaneous APs and compared the maximum rates of depolarization and repolarization. At room temperature, the maximum rate of repolarization was 76% larger in RAPNs compared to AId neurons (Fig. 2E; Table 1), whereas the maximum rate of depolarization was only 29% larger in RAPNs (Table 1). At 40°C the difference in the maximum depolarization rate disappeared, while the maximum repolarization rate was 37% greater in RAPNs (Fig. 2H; Table 1). These results suggest that the relatively slower AP repolarization is predominantly responsible for the broader AP half-width of AId neurons compared to the ultranarrow RAPN APs.

We next examined the evoked firing properties by delivering increasing, step-wise 1 sec current injections to RAPNs and AId neurons (Fig. 3A & C, examples at +500 pA). As expected, a higher firing rate was observed in both brain regions at physiological temperature compared to room temperature. In the absence of injected current, both RAPNs and AId neurons fired spontaneous APs (spontaneous APs in RAPNs also observed in extracellular slice recordings; (Wood, Lovell, Mello, & Perkel, 2011)), however, RAPNs exhibited significantly higher firing rates than AId neurons at high temperatures (Fig. 3B & D; Table 1). Moreover, we consistently observed lower firing rates in AId neurons (Fig. 3B &D), with lower instantaneous (frequency of the first two spikes) and steady state (frequency of the last two spikes) firing frequencies at all levels of current injected (Fig. 3A &C, top of each AP train). These results suggest that compared to RAPNs, the slower repolarization rate of AId neurons is likely a limiting factor for high frequency repetitive spiking.

**Figure 3.**
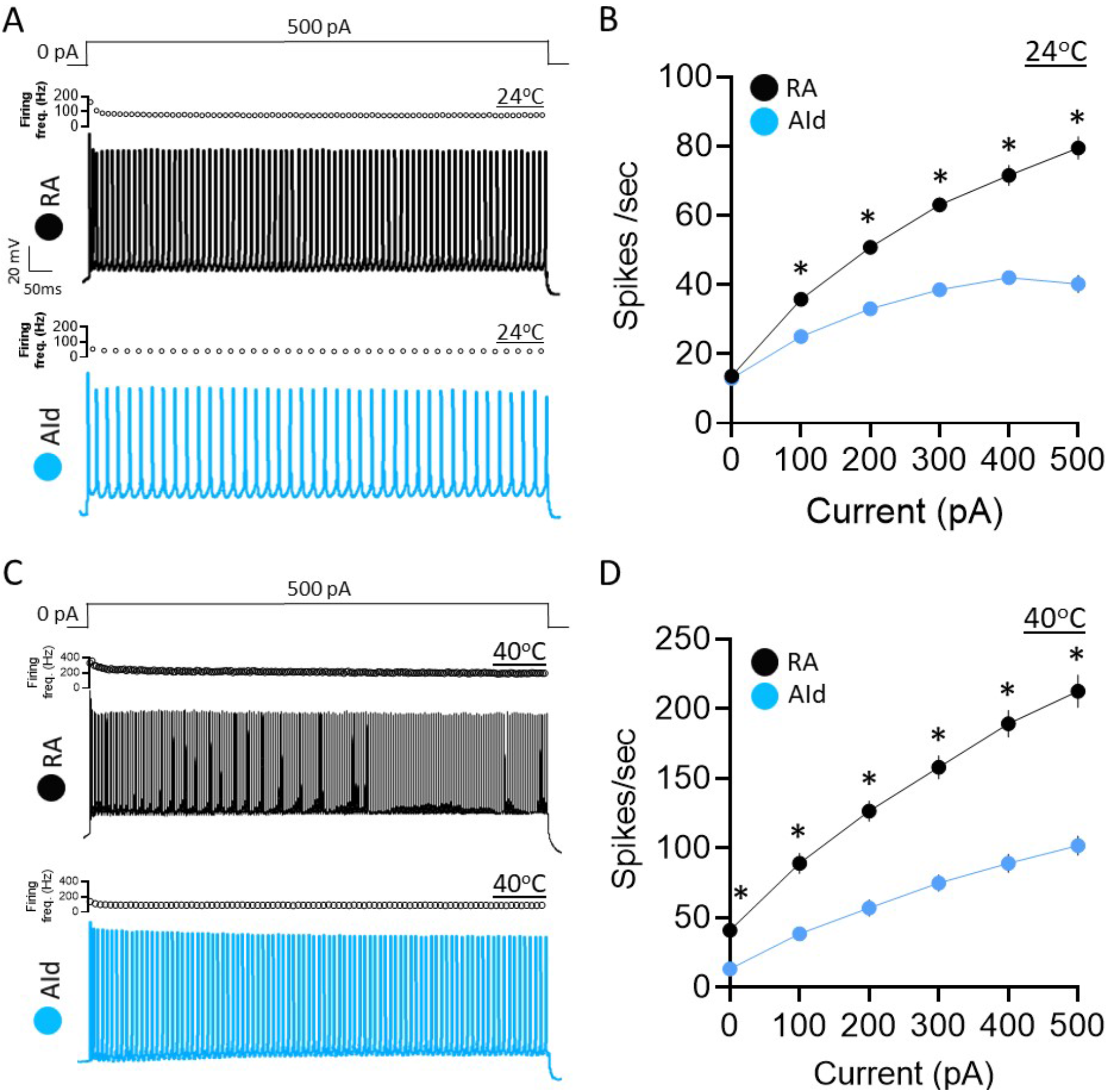
Evoked AP properties in RAPNs and AId neurons from adult male zebra finches at room and physiological temperatures. A. Representative AP trains elicited by a 1 sec 500 pA current injection at ∼24°C in a RAPN (black, top) in an AId neuron (blue, bottom). The corresponding plot of firing frequency as function of time is shown at the top of each AP train. B. Average elicited firing rate (spikes/second) as a function of injected current at ∼24°C. Two-way ANOVA; P = 5.5 × 10^−22^, F (5, 438) = 24.76, N = 39 RAPNs and 36 AId neurons. C. Representative AP trains elicited by a 1 sec 500 pA current injection at ∼40°C in a RAPN (black, top) and in an AId neuron (blue, bottom). Same scale as in (A). The corresponding plot of firing frequency as function of time is shown at the top of each AP train. D. Average elicited firing rate (spikes/second) as a function of injected current at ∼40°C. Two-way ANOVA; P =3.9 × 10^−7^, F (5, 138) = 8.69, N = 13 RAPNs and 12 AId neurons. * in C & D indicates P < 0.05; Tukey’s post-hoc test.

Within RA, inhibitory GABAergic interneurons are sparse, and are easily distinguished from RAPNs based on differences in their firing properties (Miller et al., 2017; Spiro et al., 1999; Zemel et al., 2021). *In situ* hybridization for GAD1 and 2 indicate low proportions of GABAergic cells in AId as well (zebrafinchatlas.org, (Lovell et al., 2020; Pinaud & Mello, 2007)). Accordingly, in our AId recordings we very infrequently encountered neurons with properties resembling interneurons (Supplementary Fig. 2; (Garst-Orozco et al., 2014; Kittelberger & Mooney, 1999; Liao et al., 2011; Miller et al., 2017; Spiro et al., 1999)). Because we were primarily interested in recording from excitatory neurons, and those represented the vast majority of recorded cells based on electrophysiological criteria, we excluded putative GABAergic interneurons in AId from further analysis.

### Spikes are more sensitive to Kv3 blockers in RAPNs than in AId neurons

Kv3 channels (Kv3.1-3.4) are part of the Shaw-related family of voltage gated K^+^ channels, which are characterized by rapid activation and deactivation kinetics at depolarized membrane potentials (Kaczmarek & Zhang, 2017; Rudy & McBain, 2001). These properties enable the rapid repolarization of APs, which in turn facilitates voltage gated Na^+^ (Nav) channel recovery from inactivation during repetitive firing (Bean, 2007; Leao et al., 2005; Wang et al., 1998). Kv3.1, in particular, has been implicated in narrowing the AP waveform in a number of cell types, including Betz cells (Ichinohe et al., 2004; Soares et al., 2017). Interestingly, KCNC1 (Kv3.1) mRNA is expressed within RA (Lovell et al., 2013). Moreover, RA shows much higher expression of KCNC1 than AId (Nevue et al., 2020). We thus asked whether Kv3.1 channels are regulators of the AP half-width in RAPNs and AId neurons. Although there are no commercially available Kv3.1-specific inhibitors, previous studies have established pharmacological protocols to confirm the presence of Kv3.x currents in excitable cells (Liu, Blair, & Bean, 2017; Muqeem, Ghosh, Pinto, Lepore, & Covarrubias, 2018). APs from Kv3.x-expressing neurons, including Betz cells (Spain, Schwindt, & Crill, 1991), are typically broadened by sub-millimolar concentrations of the Kv channel inhibitors tetraethylammonium (TEA) and 4-aminopyridine (4-AP) (Rettig et al., 1992). If the differences in AP half-width and maximum repolarization rate between RAPNs and AId neurons are due to the expression of Kv3.1, we would expect a more significant AP broadening upon exposure to either of these compounds in RAPNs than in AId neurons.

To test this prediction, we recorded spontaneous APs from RAPNs and AId neurons in frontal slices before and after exposure to 500 µM TEA (Fig. 4A-B) and 100 µM 4-AP (Fig. 4E-F) respectively. We performed these experiments at room temperature, as this allowed for more stable recordings that lasted for longer time periods. In response to TEA, spontaneous APs in RAPNs showed significantly more broadening (Fig. 4C; Mean ± SE: 76.7 ± 5.4% vs. 36.4 ± 7.2% average increase for RAPNs and AId neurons respectively; for separate measurements in RAPNs and AId neurons, see Supplementary Fig. 3A-B, left) and decreases in the maximum repolarization rate (Fig.4D; Mean ± SE: 48.5 ± 2.7% vs. 24.4 ± 6.2% average decrease for RAPNs and AId neurons respectively; for separate measurements in RAPNs and Aid neurons, see Supplementary Fig. 3A-B, right). In response to 4-AP, spontaneous APs in RAPNs also showed significantly more broadening (Fig. 4G; Mean ± SE: 93.0 ± 16.2% vs. 35.8 ± 9.9% average increase for RAPNs and AId neurons, respectively; for separate measurements in RAPNs and AId neurons, see Supplementary Fig. 3C-D, left) and decreases in the maximum repolarization rate (Fig. 4H; 35.1 ± 7.5% vs. 8.1 ± 7.4% average decrease for RAPNs and AId neurons, respectively; for separate measurements in RAPNs and AId neurons, see Supplementary Fig. 3C-D, right).

**Figure 4.**
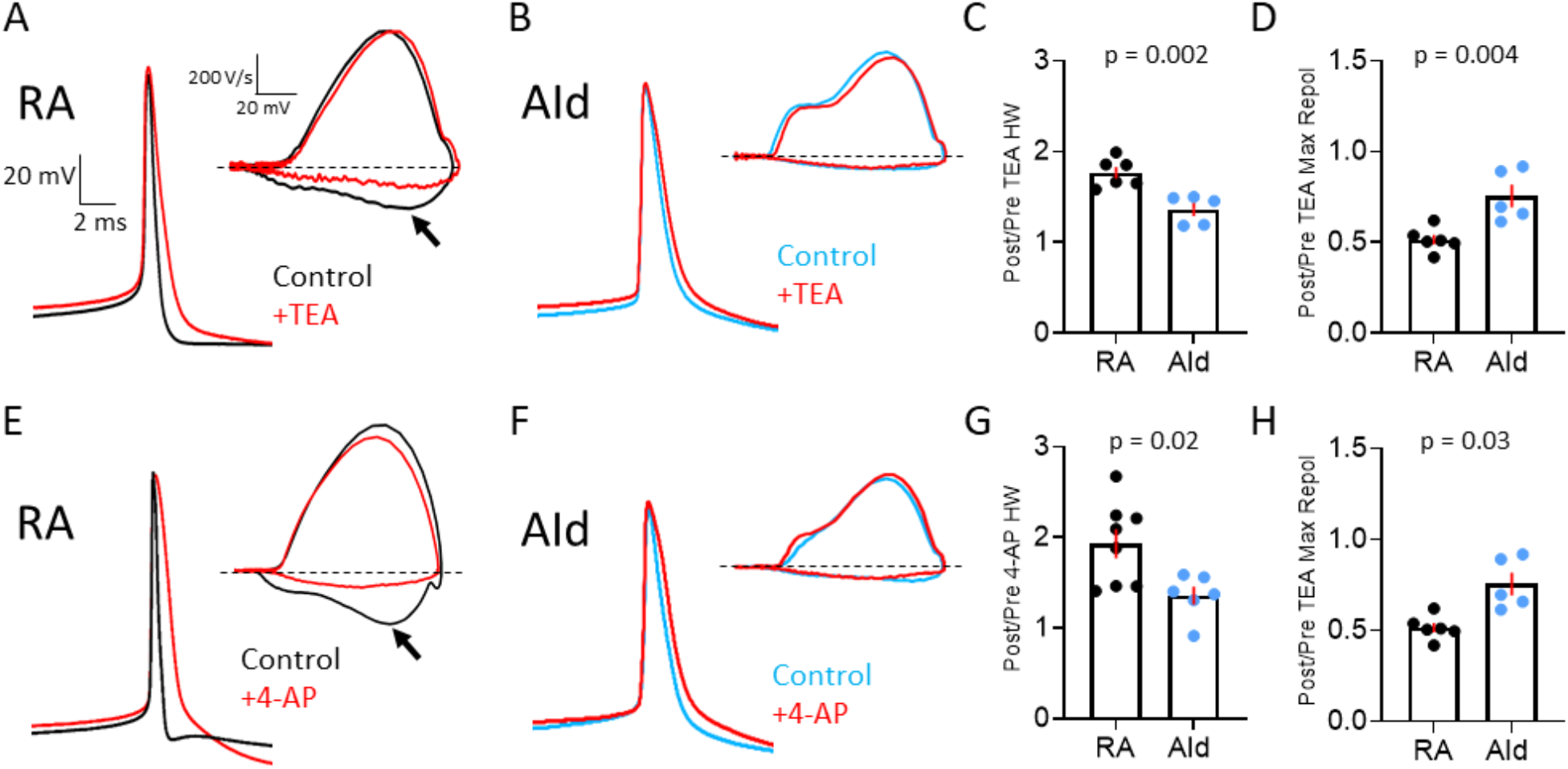
Effect of Kv3 channel inhibitors on spontaneous APs of RAPN and AId neurons. A. Representative averaged spontaneous RAPN AP traces and phase plane plots (inset) before and after 500 µM TEA administration. B. Same as in (A) with AId neurons. C. Fold change in the AP half-width (HW) in RAPNs and AId neurons (or ratio of Post/Pre TEA treatment). Student’s t-test, two-tailed, t_stat_ = 4.184, N = 6 RAPNs and 5 AId neurons. D. Fold change in the AP maximum repolarization rate in RAPNs and AId neurons. Student’s t-test, two-tailed, t_stat_ = 3.782, N = 6 RAPNs and 5 AId neurons. E. Representative averaged spontaneous RAPN AP traces and phase plane plots (inset) before and after 100 µM 4-AP administration. F. Same as in (A) with AId neurons. G. Fold change in the AP half-width in RAPNs and AId neurons. Student’s t-test, two-tailed, t_stat_ = 2.759, N = 7 RAPNs and 6 AId neurons. H. Fold change in the AP maximum repolarization rate in RAPNs and AId neurons. Student’s t-test, two-tailed, t_stat_ = 2.499, N = 7 RAPNs and 6 AId neurons. P-values are included in the graphs, red bars indicate standard error in C-D and G-H. The black arrows in A & E point to the changes in the maximum rate of repolarization. Dashed lines in A-B & E-F represent 0 V/sec.

Evoked AP firing was also more affected by TEA and 4-AP in RAPNs than in AId neurons. Upon exposure to TEA, RAPNs showed significant decreases in the evoked firing rate (Fig. 5A; 35 ± 4.1% decrease in spikes per sec, 29.3 ± 10.4% decrease in instantaneous firing and 33.7 ± 3.8% decrease in steady-state firing frequency upon the +500 pA current injection), whereas trends, but no significant effects, were seen in AId (Fig. 5B). Upon exposure to 4-AP, RAPNs also showed significant decreases in the evoked firing rate (Fig. 5C; 36.8 ± 10.9% decrease in spikes per sec, 63.7 ± 4.8% decrease in instantaneous firing frequency and 36.5 ± 12.9% decrease in steady-state firing frequency upon the +500 pA current injection), whereas trends, but no significant effects, were seen in AId (Fig. 5D).

**Figure 5.**
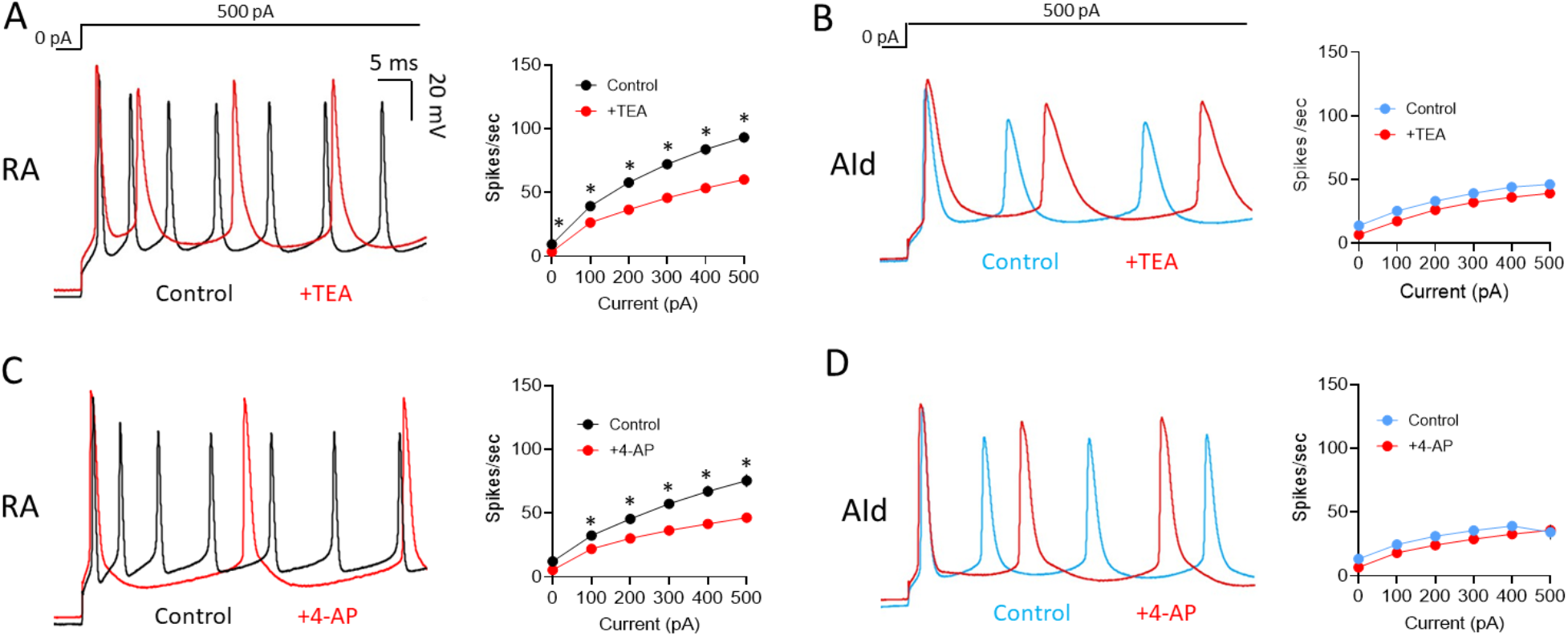
Effects of Kv3 channel inhibitors on evoked APs in RAPNs and AId neurons. A. Left to right: Representative first 50 ms of AP traces elicited by a 1 sec 500 pA current injection and average firing rates (spikes/second) before and after exposure of RAPNs to 500 µM TEA. Repeated measures two-way ANOVA; P = 6.0 × 10^−4^, F (1.279, 7.672) = 27.69, N = 7 RAPNs. B. Same as in (A) except in AId neurons. Repeated measures two-way ANOVA; P = 0.73, F (1.863, 11.18) = 0.3056, N = 7 AId neurons. C. Left to right: Representative first 50 ms of AP traces elicited by a 1 sec 500 pA current injection and average spikes/second before and after exposure of RAPNs to 100 µM 4-AP. Repeated measures two-way ANOVA; P = 8.0 × 10^−5^, F (1.256, 8.794) = 41.87, N = 8 RAPNs D. Same as in (C) except in AId neurons. Repeated measures two-way ANOVA; P = 0.26, F (1.133, 5.665) = 1.575, N = 6 AId neurons. Scales are the same in A-D. * in A-D indicates P < 0.05; Tukey’s post-hoc tests.

Our results with TEA and 4-AP alone, however, could not rule out the possible contributions of other K^+^ channels that are also hypersensitive to either TEA (Kv1, Kv7, and large-conductance Ca^2+^ -activated K^+^ (BK) channels; (Al-Sabi et al., 2010; Shen et al., 1994; Wang et al., 1998)) or 4-AP (Kv1; (Shu, Yu, Yang, & McCormick, 2007; Storm, 1988; Wu & Barish, 1992)). Thus, we next tested the effects α-dendrotoxin (DTX), XE991, and iberiotoxin (IbTX), which are highly selective antagonists of Kv1.1/1.2/1.6, Kv7, and BK channels, respectively. Upon exposing RAPNs and AId neurons to these antagonists, we observed no significant effects on spontaneous AP half-width or maximum repolarization rate, in contrast to the effects of TEA and 4-AP (Supplementary Fig. 4A-F). We also observed little to no effects on spontaneous and evoked firing frequency in RAPNs or AId neurons (Supplementary Fig. 5A-F). Thus, we conclude that Kv3.x subunits are the predominant TEA- and 4-AP-hypersensitive channels in the recorded arcopallial cells.

### RAPNs have a faster, larger, high threshold TEA-hypersensitive I_K+_ than AId neurons

To confirm that the regional differences in our current clamp recordings were indeed due to differences in the outward voltage gated K^+^ current (I_K+_), we performed voltage clamp recordings, under conditions that minimize space clamping issues (details in Methods, (Zemel et al., 2021)). After pharmacologically isolating I_K+_, we delivered sequential 200 ms test pulses to -30 mV and 0 mV, from a 5 sec holding potential at -80 mV, to preferentially activate low threshold I_K+_ or both low and high threshold I_K+_ respectively. Both RAPNs and AId neurons produced outward currents during both depolarizing steps (Fig. 6A). Consistent with a difference in expression of a high threshold, non-inactivating, I_K+_, RAPNs produced significantly larger peak outward currents 200 ms (I_200ms_) after stepping to 0 mV (Fig. 6A-B; Mean ± SE: 12.2 ± 1.2 nA vs. 4.8 ± 0.7 nA for RAPNs and AId neurons, respectively), whereas outward currents were indistinguishable between RA and AId during the -30 mV step (Fig. 6B; Mean ± SE: 0.8 ± 0.1 nA vs. 0.4 ± 0.04 nA for RAPNs and AId neurons, respectively). We also noted an initial A-type current waveform component in both RAPNs and AId neurons that preceded the larger delayed rectifier component during the 0 mV step (Fig 6A, red shaded area of the current at the 0 mV test pulse). Interestingly, the time to this peak was shorter in RAPNs than in AId neurons (Fig. 6C; Mean ± SE: 4.7 ± 0.4 ms vs. 6.2 ± 0.3 ms for RAPNs and AId neurons, respectively), even though the I_K+_ size was much larger in the RAPNs.

**Figure 6.**
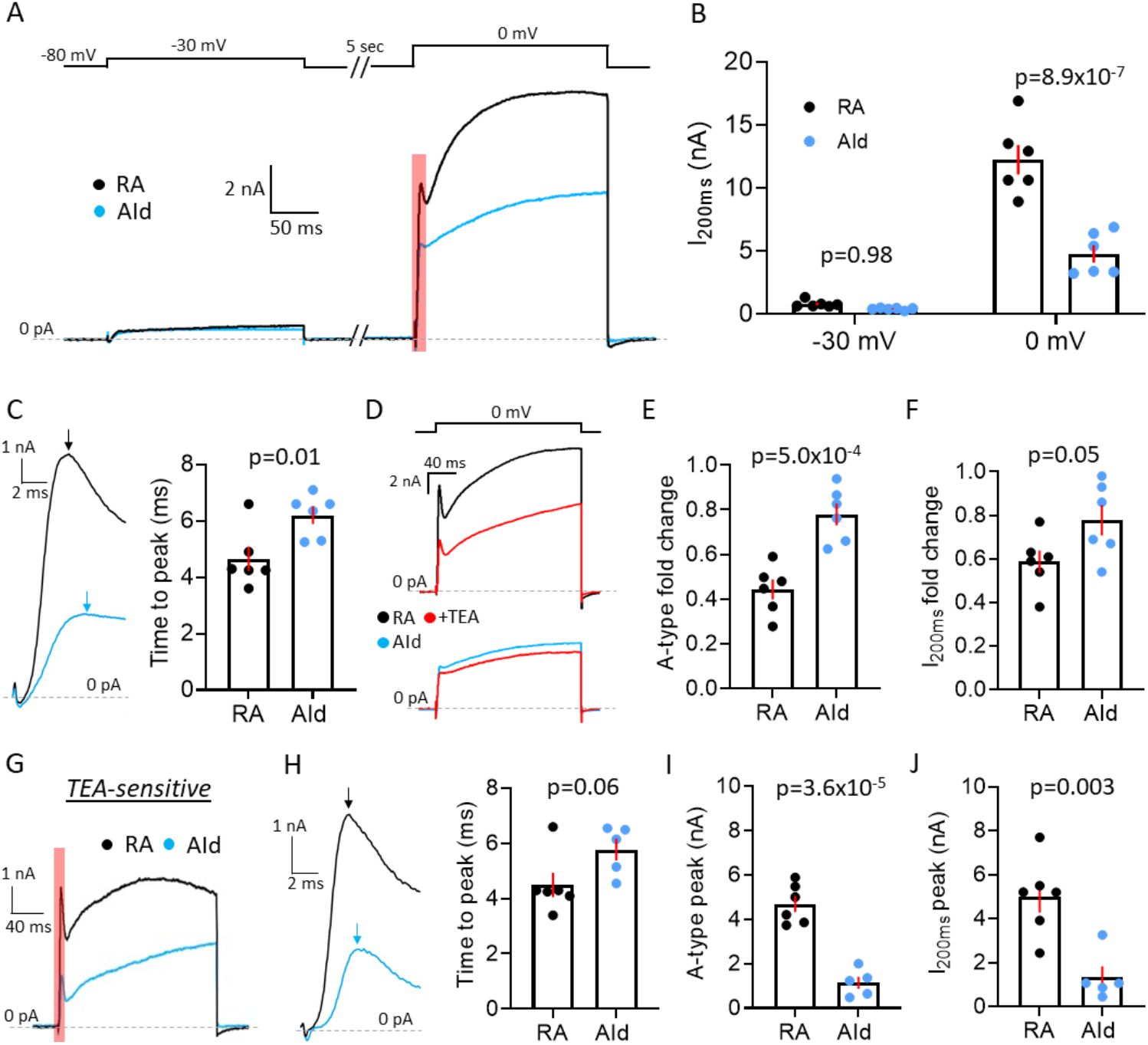
Comparison of I_K+_ between RAPNs and Aid neurons. A. Representative voltage clamp recordings of I_K+_ in a RAPN and an AId neuron at -30 mV and 0 mV. Shown at the top are the test pulses separated by 5 sec at -80 mV, with a 5 sec inter-sweep interval at -80 mV. B. Comparison of the peak current at 200 ms (I_200ms_) during both the -30 mV and 0 mV test pulses in RAPNs and AId neurons (Two-way ANOVA: P = 3.8 × 10^−5^, F (1,20) = 27.62, N = 6 RAPNs and 6 AId neurons). Red bars indicate SEM. Individual P-values determined by Tukey’s post-hoc test. C. Left: Expanded view of the A-type current component highlighted in (A). Arrows point to the peak of the A-type component of the current. Right: Comparison of the time to peak of both currents as measured from the onset of the voltage step to the peak of the A-type current (Student’s t-test, t_stat_ = 2.951, N = 6 RAPNs and 6 AId neurons). D. Representative voltage clamp recordings of a RAPN (black; top) and AId neurons (blue; bottom) before and during exposure to 500 µM TEA (red) during at the 0 mV test pulse. E. Comparison of the fold change in the initial A-type peak of I_K+_ elicited at 0 mV between RAPNs and AId neurons (Student’s t-test, two-tailed, t_stat_= 5.062, N = 6 RAPNs and 6 AId neurons). Red bars indicate SEM. F. Comparison of the fold change in the I_200ms_ of the I_K+_ elicited at 0 mV between RAPNs and AId neurons (Student’s t-test, two-tailed, t_stat_= 2.200, N = 6 RAPNs and 6 AId neurons). Red bars indicate SEM. G. Representative TEA-sensitive currents from the 0mV test pulse in an RAPN (black) and an AId neuron (blue). H. Left: Expanded view of the A-type current component highlighted in (A). Arrows pointing to the peak of the A-type component of the current highlighted in (G). Right: Comparison of the time to peak of both currents as measured from the onset of the voltage step to the peak of the A-type current (student’s t-test, t_stat_ = 2.118, N = 6 RAPNs and 5 AId neurons). I. Comparison of the peak of the A-type component of the TEA-sensitive current (student’s t-test, t_stat_ = 7.528, N = 6 RAPNs and 5 AId neurons). Red bars indicate SEM. J. Comparison of the I_200ms_ peak (student’s t-test, t_stat_ = 4.028, N = 6 RAPNs and 5 AId neurons). Red bars indicate SEM.

Informed by our current clamp pharmacology experiments, we next tested the prediction that I_K+_ would be more attenuated by sub-millimolar concentrations of TEA in RAPNs than in AId neurons. Upon washing in 500 µM TEA onto slices we saw decreases in the peak current from neurons in both brain regions (Fig. 6D), with RAPNs showing a larger fold-change in both the peak of the A-type current component (Fig. 6E; Mean ± SE: 55.7 ± 4.4% vs. 22.0 ± 4.9% decrease for RAPNs and AId neurons, respectively) and in the peak current at I_200ms_ (Fig. 6F; Mean ± SE: 41.2 ± 5.2% vs. 22.2 ± 7.0% decrease for RAPNs and AId neurons, respectively). We then extracted the TEA-sensitive I_K+_ by subtracting the post-TEA traces from the pre-TEA traces. Like the pre-TEA currents obtained at 0 mV, the TEA-sensitive current in both RAPNs and AId neurons had A-type and delayed rectifier components (Fig. 6G). By dividing the current 8 ms after the initial A-type peak by the peak A-type current we found that the degree of inactivation was indistinguishable between RAPNs and AId neurons (Supplementary Fig. 6). Notably, the time to peak of the A-type component was preserved (Mean ± SE: 4.5 ± 0.4 ms vs. Mean ± SE: 5.8 ± 0.4 ms for RAPNs and AId neurons, respectively), with RAPNs maintaining a trend toward smaller values (Fig. 6H). Importantly, whether measuring the A-type current peak (Fig. 6I; Mean ± SE: 4.7 ± 0.4 nA vs. 1.1 ± 0.3 nA for RAPNs and AId neurons, respectively) or the I_200ms_ peak (Fig. 6J; Mean ± SE: 5.0 ± 0.7 nA vs. 1.3 ± 0.5 nA for RAPNs and AId neurons, respectively), RAPNs had a significantly larger TEA-sensitive I_K+_ than AId neurons. In sum, our voltage clamp results show a higher proportion of a fast activating, TEA-hypersensitive current in RAPNs compared to AId neurons, consistent with the presence of a larger Kv3 current in RAPNs.

### Kv3.1 is the TEA-hypersensitive ion channel subunit preferentially expressed in RAPNs

The evidence presented thus far is consistent with the ultra-fast APs unique to RAPNs being related to higher expression of Kv3.1 in RA compared to AId. However, Kv3.1 is only one of four members of the Shaw-related channel family (Kv3.1-3.4) in vertebrates (Kaczmarek & Zhang, 2017; Rudy & McBain, 2001). To solidify a link between RAPN properties and Kv3.1, it was important to examine other Kv3.x family members. As a start, through close assessments of reciprocal alignments and synteny we confirmed that the locus named KCNC1 (100144433; located on chromosome 5) is the zebra finch ortholog of mammalian KCNC1, noting the conserved synteny across major vertebrate groups (Supplementary Fig. 7A). Importantly, the predicted zebra finch Kv3.1 protein (Kv3.1b isoform) is remarkably conserved (96.48% residue identity) with human (Supplementary Fig. 8) and is thus predicted to have similar pharmacology as in mammals, which is supported by our recordings from RAPNs. Additionally, we confirmed the correct identification of the zebra finch orthologs of mammalian KCNC2/Kv3.2 and KCNC4/Kv3.4, as previously reported (Lovell et al., 2013).

Previous investigations of the zebra finch genome (taeGut1, (Warren et al., 2010)) reported that the zebra finch lacked a KCNC3/Kv3.3 ortholog (Lovell et al., 2013). Recent long-read sequencing technology, however, has facilitated a more complete assembly of genomes (Rhie et al., 2021), elucidating the presence of some genes previously thought to be absent. Using the recent RefSeq release 106 (GCF_003957565.2 assembly), we observed a locus (LOC115491734) on the newly assembled zebra finch chromosome 37 described as similar to member 1 of the KCNC family. LOC115491734, however, exhibited the highest alignment scores and conserved upstream synteny with KCNC3/Kv3.3 in humans and various vertebrate lineages, noting that NAPSA in zebra finch and other songbirds is misannotated as cathepsin D-like (LOC121468878) (Supplementary Fig. 7C). We conclude that LOC115491734 is the zebra finch ortholog of mammalian KCNC3, previously thought to be missing in birds (Lovell et al., 2013), and not KCNC1. We note that the downstream immediate synteny is not conserved across vertebrate lineages, with the ancestral condition in tetrapods likely being TBC1D17 downstream of KCNC3. The predicted zebra finch KCNC3 protein showed only moderate conservation with human (68.32% residue identity), some domains including the BTB/POZ and transmembrane domain being fairly conserved, but spans of residues on the N-terminal and C-terminal regions being highly divergent. Notably, the N-terminal inactivation sequence (Rudy & McBain, 2001) of KCNC3 appears to be absent in the zebra finch (Supplementary Fig. 9).

We next performed *in situ* hybridization for all identified KCNC/Kv3.x family members in adjacent frontal brain sections from adult male zebra finches. We replicated our previous finding that Kv3.1 expression is higher in RA than in AId ((Nevue et al., 2020); Fig. 7A, top left), whereas expression of both Kv3.2 (Fig. 7A, top right) and Kv3.4 (Fig. 7A, bottom right) was non-differential between RA and AId. Unlike the graded distribution found in avian (Parameshwaran, Carr, & Perney, 2001) and mammalian (Li, Kaczmarek, & Perney, 2001) auditory brainstem, Kv3.1 appeared uniformly distributed across RA. Kv3.2-expressing cells were sparse, strongly labeled, and reminiscent of the GABAergic cell distribution (Pinaud & Mello, 2007), while Kv3.4 expression was uniformly weak throughout both brain regions (Fig. 7A). We also found that Kv3.3, while a uniquely specific marker for both brain regions compared to the surrounding arcopallium, is non-differentially expressed between RA and AId (Fig. 7A, bottom left). Importantly, we found strong Kv3.3 expression in the Purkinje cell layer of the cerebellum (Fig. 7A, bottom left), consistent with findings in mammals (Akemann & Knopfel, 2006). Furthermore, other TEA-hypersensitive potassium channel subunits examined, namely members of the Kv1 (KCNA), BK (KCNMA), and Kv7 (KCNQ) families, had similar expression in RA and AId, with the exception of KCNQ2, which had a lower proportion of labeled cells in RA than in AId (Fig. 7B,C). These results point to KCNC1 as the only TEA-hypersensitive subunit more highly expressed in RA compared to AId, providing supporting evidence for a role of Kv3.1 in shaping the ultranarrow AP of RAPNs.

**Figure 7.**
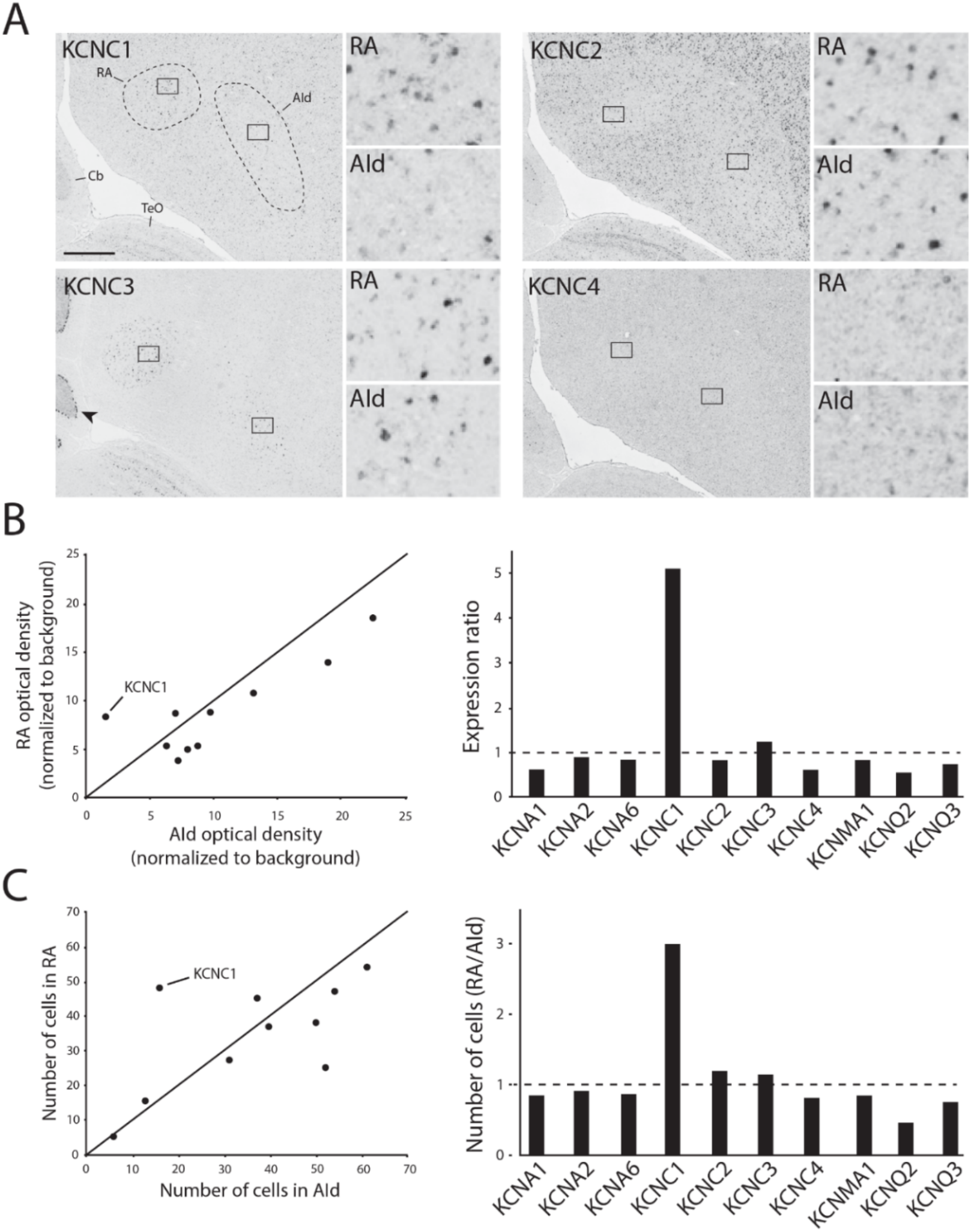
: K^+^ channel diversity in the zebra finch arcopallium: Stronger expression of KCNC1 (Kv3.1 subunit) transcripts in RA than in AId. A. Representative *in situ* hybridization images for Kv3 channel family member transcripts in RA (left) and AId (right), from nearly adjacent frontal sections of adult males. Squares in large images depict position of counting windows and of inset images for RA and AId. Arrow points to strong Kv3.3 mRNA staining in the Purkinje cell layer in the cerebellum. *Cb-* cerebellum, *TeO-* Optic tectum. Scale bar: 500 µm. B. Optical density measurements (background subtracted) in RA and AId for subunits associated with TEA-hypersensitive Kv channel types. Expression ratio (right) was calculated as RA_OD_/AId_OD_. C. Labeled cell counts in RA and AId for subunits associated with TEA-hypersensitive Kv channel types. Cell count ratios of RA/AId are shown on right.

Intriguingly, while curating avian KCNC3s, we discovered a previously undescribed KCNC family member in the genomes of several bird species, but notably absent in songbirds (Supplementary Fig. 7B). This gene most closely resembled KCNC1 in predicted domains and amino acid conservation, thus we named this KCNC1 paralog as KCNC1L. We also observed KCNC1L in non-avian sauropsids including lizards and snakes, where the locus seems to be duplicated. It was not present in humans/mammals, nor in amphibian or fish outgroups. This suggests this paralog possibly arose after the split between mammals and sauropsids, with a subsequent loss in songbirds. While helping to further solidify the differential expression of KCNC1/Kv3.1 as key for RAPN physiology, these findings bring new insights into the evolution of this important family of neuronal excitability regulators in vertebrates. How the newly identified avian KCNC3 and non-oscine KCNC1L contribute to avian neuronal physiology are intriguing questions for further study.

### AUT5 narrows the AP half-width and increases the firing rate of RAPNs

Whereas pharmacological inhibitors provided strong evidence of a major role for Kv3.x channels in RAPNs compared to AId neurons (Fig. 4-6), they notably lack the specificity for Kv3.x subunits. In contrast, the novel positive Kv3.1/3.2 modulator, AUT5, potentiates Kv3.1 currents by speeding up activation kinetics and leftward shifting the voltage dependence of activation (Taskin et al., 2015). To examine AUT5 effects, we recorded both spontaneous (Fig. 8A & C, left) and evoked (Fig. 8B & D, left) APs from RAPNs and AId neurons in the whole cell current clamp configuration, before and during exposure to 1 µM AUT5 (EC_50_ = 1.3 µM (Taskin et al., 2015)). Compared to pre-drug measurements, RAPNs showed a significant narrowing of the AP waveform (Mean ± SE: 6.5 ± 2.1% decrease; Fig. 8A, middle) and an increase of the maximum repolarization rate (Mean ± SE: 12.2 ± 4.3% increase; Fig. 8A, right) but not of the maximum depolarization rate (paired t-test, t_stat_ = 1.04, P = 0.3), whereas no significant effects were seen in AId neurons (Fig. 8C, middle and right). Spontaneous APs from RAPNs also displayed significant depolarizations in the peak (Mean ± SE: 35.1 ± 1.1 mV to 38.9 ± 1.2 mV, paired t-test, t_stat_ = 2.72, P = 0.03), threshold (Mean ± SE: -58.9 ± 1.8 to -57.0 ± 1.6, paired t-test, t_stat_ = 3.542, P = 0.01), with a non-significant trend observed for the peak afterhyperpolarization (Mean ± SE: -72.0 ± 0.7 to -71.1 ± 0.7, paired t-test, t_stat_ = 2.3, P = 0.06). In comparison AId only showed a modest change in the afterhyperpolarization (−69.6 ± 1.1 to -67.3 ± 0.6, paired t-test, t_stat_ = 2.874, P = 0.03). These changes in the spike waveform correlated with a significant increase in evoked spikes produced in RAPNs that was not seen in AId neurons (Fig. 8B & D. Mean ± SE: 25.7% ± 6.1% for RAPNs during the 1 sec +500 pA current injection). This effect is consistent with AUT5 effects observed in the Kv3.1 expressing hippocampal GABAergic interneurons of rats (Boddum et al., 2017). Interestingly, whereas AUT5 had no effects on the instantaneous firing frequency in RAPNs, the steady-state firing frequency (as measured for the last two spikes recorded during the 1 sec +500 pA current injection) increased substantially (Mean ± SE: 24.3 ± 8.2% increase). Taken together, these results are again consistent with the finding of higher Kv3.1 expression in RAPNs than in AId, and further support the role for Kv3.1 in the specialized fast firing properties of RAPNs.

**Figure 8.**
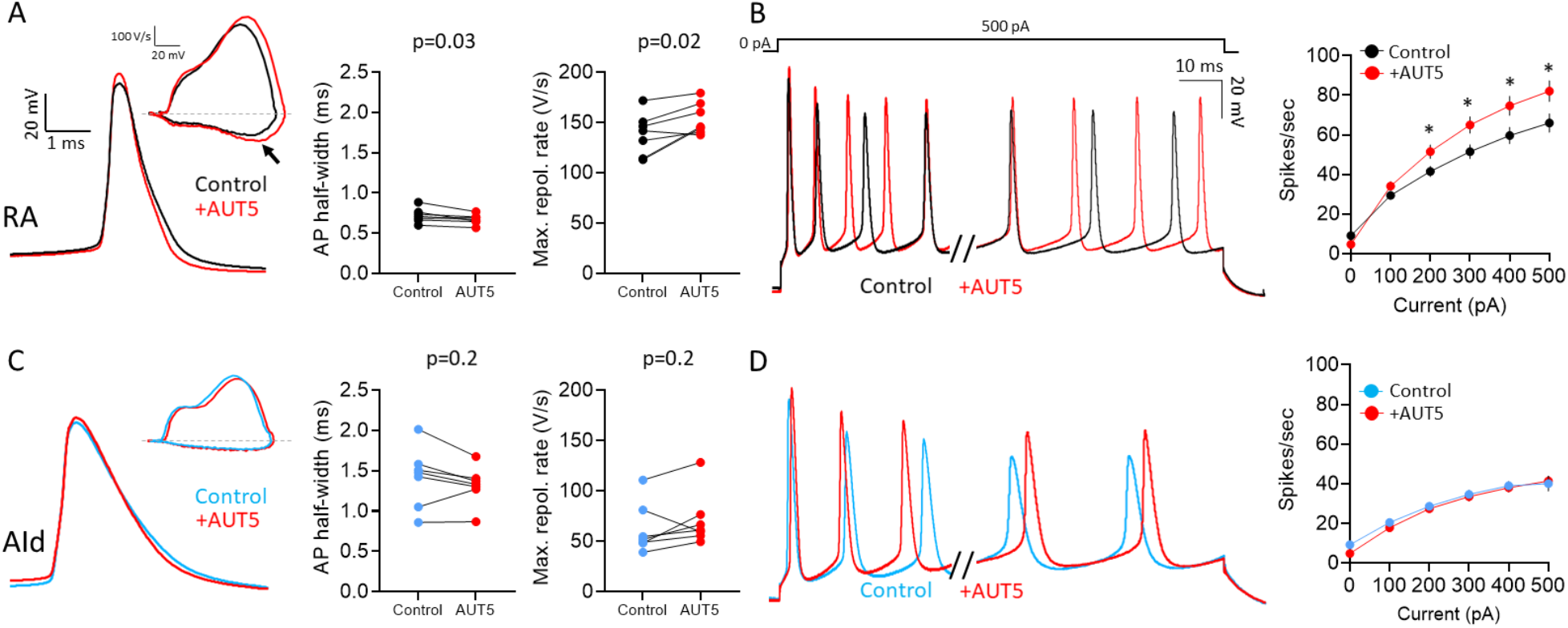
Effects of the Kv3.1/3.2 positive modulator AUT5 on APs of RAPN and AId neurons. A. Left to right: Representative AP traces, phase plane plots (inset), and changes in AP half-width and maximum repolarization rate upon exposure to 1 µM AUT5 for RAPNs. The black arrow points to the change in the maximum rate of repolarization. AP half-width: Paired student’s t-test, two-tailed, t_stat_ = 2.838, N= 7 RAPNs; Maximum repolarization rate: Paired student’s t-test, two-tailed, t_stat_ = 3.095, N= 7 RAPNs. B. Left to right: First 40 ms and last 65 ms of representative AP traces elicited by a 1 sec 500 pA current injection and average evoked firing rates (spikes/second) before and after exposure to 1 µM AUT5 for RAPNs. Repeated measures two-way ANOVA; P = 3.1×10^−12^, F (5, 30) = 32.40, N = 7 RAPNs; * indicates P < 0.05; Tukey’s post-hoc tests. C. Left to right: Representative AP traces, phase plane plots (inset), and changes in AP half-width and maximum repolarization rate upon exposure to 1 µM AUT5 for AId neurons. AP half-width: Paired student’s t-test, t_stat_ = 1.483, N= 7 AId neurons, two-tailed; Maximum repolarization rate: Paired student’s t-test, two-tailed, t_stat_ = 1.523, N= 7 AId neurons. Same scale as in (A). D. Left to right: First 40 ms and last 65 ms of representative AP traces elicited by a 1 sec 500 pA current injection and average evoked firing rates (spikes/second) before and after exposure to 1 µM AUT5 for AId neurons. Repeated measures two-way ANOVA; P = 0.18, F (5, 30) = 1.658, N = 7 AId neurons. Same scale as in (B).

## Discussion

Our results demonstrate that RAPNs and AId neurons exhibit some common properties, including spontaneous firing, non-adapting firing during current injections and similar expression profiles of KCNC2-4 (Kv3.2-3.4) subunits. However, they differ greatly in AP spike half-width and capacity for high-frequency firing. We show that these unique RAPN properties, which are reminiscent of the large Betz cells in layer 5 of primate M1 cortex, are associated with higher expression of KCNC1 (Kv3.1) in RA than in AId. In contrast, the properties of AId neurons, which are involved in non-vocal somatic motor function in birds, are more similar to those of canonical mammalian L5PNs. We propose that RAPNs are highly specialized neurons for song production, sharing unique functional and molecular properties with primate Betz cells. This allows both cell types to reliably fire ultranarrow spikes at high frequencies.

### Songbird upper motor neurons have distinct morphologies

The morphology of RAPNs described here is consistent with previous morphological characterizations, suggesting that these neurons represent a fairly homogeneous cell type (Kittelberger & Mooney, 1999; Spiro et al., 1999). In contrast, this study includes a first attempt to describe morphological features of AId neurons (Fig. 1; Supplementary Fig. 1). We found that RAPNs and AId neurons share similar soma size and elaborate branched dendrites, and both form extensive local axon collaterals. However, we also obtained evidence of morphological differences that may have implications for how RAPNs and AId neurons integrate synaptic inputs. The higher dendritic branch complexity revealed by a Sholl analysis (D. A. Sholl, 1956) and the higher number of dendritic spines (Fig. 1A; Supplementary Fig. 1C) of AId neurons (a finding revealed using two methods: FIJI (Schindelin et al., 2012) and ShuTu (Jin et al., 2019)), suggests that AId may receive more synaptic inputs than RAPNs. Such inputs would be from the nidopallium, the main known source of input to the AI (Bottjer et al., 2000; Johnson, Sablan, & Bottjer, 1995; Karten, 2015), and/or from local GABAergic interneurons. Conversely, the higher proportion of large spines in RAPNs than in AId neurons (Supplementary Fig. 1D) parallels Betz cells, which also contain higher proportions of large spines compared to other pyramidal neurons within the cat M1 (Kaiserman-Abramof & Peters, 1972). These differences in spine shape and surface area suggest potential differences in filtering of incoming synaptic signals, with some input sources potentially having larger effects on membrane potential changes than others. Of note, the difference spine density between RAPNs and AId closely approximate findings in the high vocal center (HVC), where upstream HVC-RA projecting neurons exhibit roughly half the spine density as HVC-Area X projecting neurons (Kornfeld et al., 2017). Importantly, whereas axonal projection targets for RAPNs are discrete and have been studied in detail (Vicario, 1991; Wild, 1993b; Wild, Kubke, & Mooney, 2009; Wild, Williams, & Suthers, 2001), those for AId neurons appear to be more complex (Bottjer et al., 2000) and are possibly more heterogeneous in terms of cell type composition. While our present findings do not address potential subpopulations of AId neurons, they lay the groundwork for future efforts using cell filling and/or track tracing to further characterize the similarities and differences between RAPNs and AId neurons.

Calculations of membrane capacitance (C_m_) allowed us to estimate the average surface area of RAPNs and AId neurons (Table 1), whereas 3-D reconstructions using ShuTu allowed us to also estimate shrinkage-corrected surface areas (Supplementary Table 1). The values were fairly close, in spite of the uncertainties and assumptions involved in both methods (see Results).

Based on their larger overall spine surface area, it appears that AId neurons are investing more “resources” on spines than RAPNs, despite having proportionately smaller individual spines. We suggest that the sparse population of large spines in adult male RAPNs may limit filtering of synaptic inputs compared to AId neurons, a factor that could at least partly contribute to the highly stereotyped song of adult male finches. Large, stable and mature spines in RAPNs may also enable large EPSCs currents in adult RAPNs from male birds with crystallized songs (Garst-Orozco et al., 2014; Kittelberger & Mooney, 1999). Future electrophysiology and morphology studies are needed to address this question further.

### Shared properties between upper-motor neuron subclasses in finches and mammals

Comparative studies in the auditory system have suggested convergent strategies for temporal coding of sound stimuli in birds and mammals (Carr & Soares, 2002; Spool et al., 2021). We have recently described how upper-motor RAPNs and L5PNs in mammalian M1 also share convergence in factors determining the AP initiation and upstroke. This includes APs with biphasic depolarization rates, persistent Na^+^ currents, large transient Na^+^ currents (I_NaT_) with rapid kinetics, and high Navβ4 mRNA expression, which we showed to be linked to robust resurgent currents (I_NaR_) in RAPNs (Zemel et al., 2021). A major difference, however, was the ultranarrow AP waveform of RAPNs that is also a unique trait of the large Betz cells found in Layer 5 of M1 in cats (Chen et al., 1996) and primates (Vigneswaran et al., 2011). Here we show RAPNs produce narrower APs compared to AId neurons, due largely to higher maximum repolarization rates (Fig. 2). In contrast to RAPNs, the 1.2 ms AP half-width at 24°C and the “regular” non-adapting AP firing of AId neurons render them more similar to canonical L5PNs in the motor cortex of rodents (Lacey et al., 2014), cats (Chen et al., 1996) and primates (Vigneswaran et al., 2011). Thus, like Betz cells (Bakken et al., 2021; Ichinohe et al., 2004; Soares et al., 2017), RAPNs appear to be a specialized class of upper motor neurons that display higher temporal precision of AP firing and faster firing rates than AId neurons.

We also note striking differences between the properties of finch RAPNs and AId neurons compared to those of L5PNs in mammalian M1. Foremost, RAPNs and AId neurons lack the large, tufted apical dendrites typical of L5PNs (Network, 2021). Additionally, these avian neurons fire spontaneously in the absence of synaptic inputs, a property not typically seen in M1 L5PNs (Chen et al., 1996; Lacey et al., 2014). The size and distribution of Betz cells in M1 also differ compared to RAPNs in the finch. Whereas Betz cells have very large somas and are interspersed with smaller L5PNs (Lassek, 1940; Rivara et al., 2003), RAPNs and AId neurons have similar soma sizes and localize to adjacent but distinct regions within the finch arcopallium (Bottjer et al., 2000; Johnson et al., 1995; Nevue et al., 2020).

### Kv3.1 subunits facilitate fast spiking in RAPNs

By quickly activating at depolarized voltages, Kv3 channels can efficiently initiate Nav channel recovery from inactivation during repetitive AP firing (Bean, 2007; Gu et al., 2018; Kaczmarek & Zhang, 2017; Rudy & McBain, 2001). Importantly, the fact that the AP waveforms and firing rates of RAPNs were indeed more sensitive to sub-millimolar concentrations of TEA and 4-AP than AId neurons (Fig. 4-6) implicate Kv3.1 channels in regulating the excitable features of RAPNs. Notably, however, these antagonists have known effects on other members of this ion channel family, even at sub-millimolar concentrations (Kaczmarek & Zhang, 2017). *In situ* hybridization showed higher expression of the Kv3.1 subunit in RAPNs than in AId neurons (Fig. 7), suggesting a unique role of Kv3.1 in RAPNs. Using the Kv3.1/3.2 positive modulator AUT5, we were able to further correlate the differential expression of Kv3.1 subunits with excitable properties in RAPNs and AId neurons. AUT5 is known to increase activation kinetics, decrease deactivation kinetics, and leftward shift the voltage dependence of activation of Kv3.1 channels (Boddum et al., 2017; Taskin et al., 2015). The narrowing of the AP waveform and increase in steady state firing provide strong support for a role of Kv3.1 in the repolarization of APs and in increasing the availability of Nav channels during high frequency firing in RAPNs.

We also noted a modest depolarization of the AP peak, after-hyperpolarization and threshold with AUT5 exposure. Considering this combination of changes was not observed in AId, this may be an additional result of increases in Nav channel availability due to positive modulation of Kv3.1. The fact that the AUT5 effects on Kv3.1-expressing zebra finch neurons were similar to those seen in mammals (Boddum et al., 2017) is not surprising, as the predicted peptide sequence of finch Kv3.1 is ∼96.5% identical to that of the human Kv3.1b splice variant (Supplementary Fig. 8). Songbirds and mammals diverged 300 million years ago, so this remarkable conservation suggests that Kv3.1 channels may be optimized for enabling ultranarrow AP waveforms and high frequency firing. Accordingly, loss of function mutations in the human Kv3.1 gene result in myoclonus epilepsy and ataxia (MEAK), a disease that amongst other symptoms, presents with severe motor deficits (Barot, Margiotta, Nei, & Skidmore, 2020; Muona et al., 2015).

Kv3.1 channels have been found in fast-spiking, parvalbumin-expressing interneurons and in Layer 5 Betz cells of primate motor cortex (Bakken et al., 2021; Soares et al., 2017). The two splice variants described in mammals, Kv3.1a and 3.1b, have differing trafficking patterns and protein-protein interactions (Kaczmarek & Zhang, 2017; Rudy et al., 1999). The longer C-terminal domain of Kv3.1b appears to enable trafficking out of the soma into the axon, while providing protein kinase C phosphorylation sites that decrease the open probability of the channel. We note that in previous transcriptome sequencing efforts in the finch, five Kv3.1 transcripts were predicted (NCBI GeneID:100144433), including Kv3.1b, that contain C-terminal domains with varying lengths. Thus, there is a strong likelihood that, like mammals, RAPNs express different Kv3.1 splice variants that are differentially trafficked and/or phosphorylated based on the specific characteristics of their C-terminal domains.

In contrast to Kv3.1, Kv3.2-3.4 are non-differential between RA and AId. Unlike Kv3.1 and Kv3.2, Kv3.4 exhibits rapid inactivation at depolarized voltages (Kaczmarek & Zhang, 2017; Rudy & McBain, 2001). Interestingly, whereas in mammals Kv3.3 channels exhibit inactivation, albeit on a slower timeframe than Kv3.4 (Kaczmarek & Zhang, 2017; Rudy & McBain, 2001), the predicted finch Kv3.3 peptide lacks an N-terminal “ball-and-chain” domain thought to be associated with inactivation (Supplementary Fig. 9; (Rudy et al., 1999)). This would suggest that Kv3.4 is responsible for the small, inactivating component of TEA-hypersensitive currents measured in both RA and AId (Figure 6; Supplementary Fig. 6), possibly by participating in a hetero-multimeric complex with non-inactivating Kv3 subunits (Rudy & McBain, 2001). Interestingly, cell-attached recordings in rat layer V pyramidal neurons in the sensorimotor cortex display TEA-insensitive A-type currents, a finding consistent with our data (Kang, Huguenard, & Prince, 2000). This current is likely composed of channels that, in contrast to Kv3.4, gate at significantly more negative membrane potentials (Kang, Huguenard, & Prince, 2000). Despite lacking an inactivation domain, Kv3.3 expression in both RA and AId likely contributes to the TEA-hypersensitivity seen in both current- and voltage-clamp recordings. Interestingly, at the calyx of Held nerve terminal, which can spike at 1 kHz (Kim, Renden, & von Gersdorff, 2013), the Kv3.3 subunit disproportionately contributes to the AP waveform compared to Kv3.1 (Richardson et al., 2022). These two subunits may be differentially trafficked to subcellular compartments, including dendrites, which may facilitate burst-pause time coding (Deng et al., 2005; Zang & De Schutter, 2021).

### The combination of Kv3.1 and Navβ4 facilitates narrow and energetically efficient spikes

Birds and mammals are warm-blooded and their physiological temperatures facilitate narrow AP waveforms in a number of cell types by limiting the overlap of fast Na^+^ and K^+^ conductances, thus making the neurons more energetically efficient (Alle, Roth, & Geiger, 2009; Fohlmeister, 2009; Hu, Roth, Vandael, & Jonas, 2018). RAPNs fire spontaneously with high-frequency bursts of spikes just before and during song production (Daliparthi et al., 2019; Olveczky, Otchy, Goldberg, Aronov, & Fee, 2011; Yu & Margoliash, 1996). This is presumably an energetically costly process and, accordingly, RA in adult males displays a dense staining for cytochrome oxidase (Adret & Margoliash, 2002).

The combination of Navβ4, and its associated I_NaR_, and Kv3 channels likely promote ultranarrow AP waveforms and rapid bursting in RAPNs (Zemel et al., 2021) and fast spiking nerve terminals (Kim, Kushmerick, & von Gersdorff, 2010; Richardson et al., 2022). Studies in rodent cerebellar Purkinje (Akemann & Knopfel, 2006) and vestibular nucleus neurons (Gittis, Moghadam, & du Lac, 2010) suggest that the repolarization enabled by Kv3 currents enhances the activation of post-spike I_NaR_, likely facilitating the high frequency firing that occurs during *in vivo* bursting (Loewenstein et al., 2005; Saito & Ozawa, 2007). This is consistent for the recently identified role for I_NaR_ in stabilizing burst duration and intervals in neurons of the mesencephalic nucleus in mice (Venugopal et al., 2019). This exquisite coordination between Navs, Kvs and auxiliary subunits is likely part a broader cohort of molecular markers identified as unique to fast spiking neurons (Kodama et al., 2020; Network, 2021). Interestingly, arcopallial neurons outside of RA, and RAPNs from juvenile male finches lack Navβ4 and Kv3.1 (unpublished observations) expression, exhibit much broader APs and are incapable of high frequency firing (Zemel et al., 2021). Our results thus point to important synergistic roles of joint Navβ4 and Kv3.1 expression in shaping RAPN excitable properties.

### Similarities between RAPNs and AId neurons and the evolutionary origins of RA

The AP waveforms in RAPNs and AId neurons are similar in several parameters, including threshold, maximum depolarization rate, amplitude, peak and after-hyperpolarization (Table 1). These cells additionally have a multitude of common molecular correlates of excitability (*e*.*g*. high expression of Nav1.1, Nav1.6 and Navβ4 (Mello, Kaser, Buckner, Wirthlin, & Lovell, 2019; Nevue et al., 2020)) that likely underlie some of their physiological similarities. We have now found that both RAPNs and AId neurons express Kv3.3 subunit transcripts, a previously unrecognized gene in the songbird genome, which likely imparts some of the TEA- and 4-AP-hypersensitivity of AP waveforms. These shared properties of RAPNs and AId neurons distinguish them from those examined in other arcopallial regions (e.g. caudal arcopallium outside of RA), where neurons exhibit broad APs, are incapable of sustained fast spiking, and exhibit less expression of the aforementioned ion channel subunits (Friedrich, Lovell, Kaser, & Mello, 2019; Lovell et al., 2013; Nevue et al., 2020; Zemel et al., 2021). Coupled with evidence of AId’s involvement in somatic motor control (Feenders et al., 2008; Yuan & Bottjer, 2020), the current molecular and electrophysiological data further suggests that these two regions may share a common evolutionary origin, and that RA may have evolved as a specialized expansion of AId (Feenders et al., 2008).

In conclusion, RAPNs in zebra finches exhibit many fundamental molecular and functional similarities to primate Betz cells, that may be involved is fine digit movements (Tomasevic et al., 2022). In combination with their well-defined role in singing behaviors, this study identifies RAPNs as a novel and more accessible model for studying the properties of Betz-like pyramidal neurons that offer the temporal precision required to control complex learned motor behaviors.

## Methods

### Animal subjects

All of the work described in this study was approved OHSU’s Institutional Animal Care and Use Committee (Protocol #: IP0000146) and is in accordance with NIH guidelines.

Zebra finches (*Taeniopygia guttata*) were obtained from our own breeding colony. All birds used were male and > 120 days post hatch. Birds were sacrificed by decapitation and their brains removed. For electrophysiology experiments brains were bisected along the midline, immersed in ice-cold cutting solution, and processed as described below. For *in situ* hybridization experiments brains were cut anterior to the tectum and placed in a plastic mold, covered with ice-cold Tissue-Tek OCT (Sakura-Finetek; Torrance, CA), and frozen in a dry ice/isopropanol slurry and processed as described below.

### *In situ* hybridization

To compare mRNA expression levels for *KCNC1, KCNC2, KCNC3, KCNC4, KCNMA1, KCNQ2, KCNQ3, KCNA1, KCNA2, and KCNA6* across RA and AId, brains sections (thickness = 10 μm) were cut coronally on a cryostat and mounted onto glass microscope slides (Superfrost plus; Fisher Scientific, Hampton, NH, USA), briefly fixed, and stored at -80°C. For each brain, every 10^th^ slide was fixed and stained for Nissl using an established cresyl violet protocol. Slides were examined under a bright-field microscope to identify sections containing the core region of RA and AId as previously defined (Nevue et al., 2020). *In situ* hybridization was conducted using an established protocol (Carleton et al., 2014). Briefly, slides were hybridized under pre-optimized conditions with DIG-labeled riboprobes synthesized from BSSHII-digested cDNA clones obtained from the ESTIMA: Songbird clone collection (Replogle et al., 2008). Specific clones corresponded to GenBank IDs CK302978 (KCNC1; Kv3.1), DV951094 (KCNC2; Kv3.2), DV953393 (KCNC3; Kv3.3), CK308792 (KCNC4; Kv3.4), DV954467 (KCNMA1; BK), FE737967 (KCNA1; Kv1.1), FE720882 (KCNA2; Kv1.2), FE733881 (KCNA6; Kv1.6), DV954380 (KCNQ2; Kv7.2), and CK316820 (KCNQ3, Kv7.2). After overnight hybridization, slides were washed, blocked, incubated with alkaline phosphatase conjugated anti-DIG antibody (1:600; Roche, Basal, Switzerland) and developed overnight in BCIP/NBT chromogen (Perkin Elmer; Waltham, MA, USA). Slides were coverslipped with VectaMount (Vector, Newark, CA, USA) permanent mounting medium, and then digitally photographed at 10X under bright field illumination with a Lumina HR camera mounted on a Nikon E600 microscope using standardized filter and camera settings. Images were stored as TIFF files and analyzed further using the FIJI distribution of ImageJ (Schindelin et al., 2012). We note that high-resolution parasagittal images depicting expression of *KCNC1, KCNC2, KCNA1, KCNA6, KCNQ2*, and *KCNMA1* in RA of adult male zebra finches are available on the Zebra Finch Expression Brain Expression Atlas (ZEBrA; www.zebrafinchatlas.org). All probes were evaluated for specificity by examining their alignment to the zebra finch genome and avoiding probes with significant cross alignments to other loci (as detailed previously (Lovell et al., 2020)).

For each gene, we quantified both expression levels based on labeling intensity (i.e. average pixel intensity) and the number of cells expressing mRNA per unit area. We measured the average pixel intensity (scale: 0-256) in a 200 × 200 µm window placed over each target area in the images of hybridized sections. To normalize signal from background we subtracted an average background level measured over an adjacent control area in the intermediate arcopallium that was deemed to have no mRNA expression. The expression ratio was calculated as RA_OD_/AId_OD_ where values greater than 1 are more highly expressed in RA and values less than 1 are more highly expressed in AId. We also quantified the number of labeled cells in each arcopallial region by first establishing a threshold of expression 2.5X above the background level. Standard binary filters were applied and the FIJI ‘Analyze Particles’ algorithm was used to count the number of labeled cells per 200 µm^2^.

### Slice preparation for electrophysiology experiments

Frontal (180 μm for current clamp and 150 μm for voltage clamp) slices were cut on a vibratome slicer (VT1000, Leica) in an ice-cold cutting solution containing (in mM): 119 NaCl, 2.5 KCl, 8 MgSO_4_, 16.2 NaHCO_3_, 10 HEPES, 1 NaH_2_PO_4_, 0.5 CaCl_2_, 11 D-Glucose, 35 Sucrose pH 7.3-7.4 when bubbled with carbogen (95% O_2_, 5% CO_2_; osmolarity ∼330-340 mOsm). Slices were then transferred to an incubation chamber containing artificial cerebral spinal fluid (aCSF) with (in mM): 119 NaCl, 2.5 KCl, 1.3 MgSO_4_, 26.2 NaHCO_3_, 1 NaH_2_PO_4_, 1.5 CaCl_2_, 11 D-Glucose, 35 Sucrose pH 7.3-7.4 when bubbled with carbogen (95% O_2_, 5% CO_2_; osmolarity ∼330-340 mOsm) for 10 min at 37°C, followed by a room temperature incubation for ∼30 min prior to start of electrophysiology experiments.

### Patch clamp electrophysiology

RA and AId could be readily visualized in via infra-red differential interference contrast microscopy (IR-DIC) (Fig. 2B). Whole-cell patch-clamp recordings were performed at room temperature (∼24°C) unless otherwise indicated. For experiments performed at 40°C, the bath solution was warmed using an in-line heater (Warner Instruments, Hamden, CT). The temperature for these experiments varied up to ± 2°C.

Slices were perfused with carbogen-bubbled aCSF (1-2ml/min) and neurons were visualized with an IR-DIC microscope (Zeiss Examiner.A1) under a 40x water immersion lens coupled to a CCD camera (Q-Click; Q-imaging, Surrey, BC, Canada). Whole-cell voltage- and current-clamp recordings were made using a HEKA EPC-10/2 amplifier controlled by Patchmaster software (HEKA, Ludwigshafen/Rhein, Germany). Data were acquired at 100 kHz and low-pass filtered at 2.9 kHz. Patch pipettes were pulled from standard borosilicate capillary glass (WPI, Sarasota, FL, USA) with a P97 puller (Sutter Instruments, Novato, CA). All recording pipettes had a 3.0 to 6.0 MΩ open-tip resistance in the bath solution. Electrophysiology data were analyzed off-line using custom written routines in IGOR Pro (WaveMetrics, Lake Oswego, OR, USA).

For current clamp recordings, intracellular solutions contained (in mM): 142.5 K-Gluconate, 21.9 KCl, 5.5 Na_2_-phosphocreatine, 10.9 HEPES, 5.5 EGTA, 4.2 Mg-ATP and 0.545 GTP, pH adjusted to 7.3 with KOH, ∼330-340 mOsm. Synaptic currents were blocked by bath applying Picrotoxin (100 μM), DL-APV (100 μM), and CNQX (10 μM) (Tocris Bioscience) for ∼3 min prior to all recordings. To initiate current clamp recordings, we first established a giga-ohm seal in the voltage clamp configuration, set the pipette capacitance compensation (C-fast), and then set the voltage command to -70 mV. We then applied negative pressure to break into the cell. Once stable, we switched to the current clamp configuration. Experiments in current clamp were carried out within a 15-min period. AP half-width was defined as the width of the AP half-way between threshold (when the rate of depolarization reaches 10 V/s) and the AP peak. The maximum depolarization and repolarization rates were obtained from phase plane plots generated from averaged spontaneous APs. We noted that the resting membrane potential tended to hyperpolarize to the same degree (∼10 mV) in both RA and AId after positive current injections during these current clamp recordings (Alexander, Mitry, Sareen, Khadra, & Bowie, 2019). Recordings in which the resting membrane potential deviated by > 10 mV were discarded. We note that recordings were not corrected for a calculated liquid junction potential of +9 mV.

Estimated current clamp measurements of membrane capacitance (C_m_) were calculated from the measured membrane time constant (τ_m_; fit with a double exponential at the onset of a negative current injection) and input resistance (R_in_; calculated slope of V-I plot) using the equation:

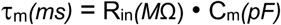

For voltage clamp recordings we attempted to limit the speed and space clamp error by 1) Cutting thinner slices (∼150 µm) to eliminate more processes, 2) decreased the intracellular K^+^ concentration to decrease the driving force and 3) compensated the series resistance to 1MΩ. The intracellular solutions contained the following (in mM): 75 K-Gluconate, 5.5 Na_2_-phosphocreatine, 10.9 HEPES, 5.5 EGTA, 4.2 Mg-ATP, and 0.545 GTP, pH adjusted to 7.3 with KOH, adjusted to ∼330-340 mOsm with Sucrose. R_s_ was compensated to 1 MΩ, the uncompensated R_s_ = 7.7 ± 0.4 MΩ (Mean ± SE; N=12 RAPNs and AId neurons). We ensured exclusion of interneurons by briefly observing the AP waveform in current clamp prior to switching to voltage clamp (Supplementary Fig. 2; (Zemel et al., 2021)). In order to isolate K^+^ currents, slices were exposed to bath applied CdCl_2_ (100 μM), TTX (1 μM), Picrotoxin (100 μM), CNQX (10 μM), and APV (100 μM) for ∼5 min prior to running voltage clamp protocols. After protocols were applied, TEA (500 μM) was bath applied and the same voltage-clamp protocols were repeated after K^+^ currents were eliminated. K^+^ currents were isolated by subtracting the TEA-insensitive current traces from the initial traces. Capacitive currents generated during voltage-clamp recordings were eliminated by P/4 subtraction. Recordings were not corrected for a measured liquid junction potential of +12 mV.

### Morphology

For these experiments Biocytin (4mg/ml; Sigma, St. Louis, MO, USA) and Alexafluor 488 (Life Technologies, Carlsbad, CA, USA) were included in the intracellular solution used in current clamp experiments. Upon entering the whole cell current clamp configuration, the current was set to 0 pA and the cell was held in for 20 to 30 minutes at room temperature to allow for complete filling. The electrode was then slowly removed at a diagonal angle as the fluorescence was monitored to determine when the electrode detached. Upon complete separation of the electrode from the cell, the slice was placed in 4% paraformaldehyde overnight. The slice was then washed in PBS with 0.25% Triton-100x (3x for 10 minutes), blocked in a PBS solution containing 1% skim milk (1 hr) and then stained in a PBS solution containing 1% skim milk and avidin conjugated to Alexafluor 594 (1:200, Life Technologies, Carlsbad, CA, USA) for 2 hrs. The slice was then washed in PBS with 0.25% Triton-100x (3x for 10 minutes) before a final wash in PBS followed by mounting on a cover slip. Images were captured with a Zeiss LSM 980 Airyscan 2 confocal microscope.

Maximum projections of the images were made and analyzed in FIJI-Image J (Schindelin et al., 2012) to calculate the 2-D soma area, dendritic complexity (D. A. Sholl, 1956), and estimates spine density (as measured from averaged 30 µm stretches from multiple tertiary branches in each cell) shown in Fig. 1. In a subset of these neurons (3 RAPNs and 2 AId neurons) we used the recently developed, open-source software, ShuTu (Jin et al., 2019), to reconstruct the morphologies in 3-D (see examples in the Supplementary Fig. 1 A-B; Video1-2). 3-D renderings were expanded 2-fold in the Z-direction to allow for better visualization of cellular processes. Despite the planar appearance of reconstructed neurons, previous imaging work in RAPNs and hippocampal neurons suggest these dendrites tile equally across planes (Spiro et al., 1999) and have been disproportionately compressed in the z-axis as a result of tissue processing (Pyapali, Sik, Penttonen, Buzsaki, & Turner, 1998) respectively. We used ShuTu’s automatic reconstruction script to trace high-contrast neurites filled with biocytin and to estimate their diameters. For low-contrast, broken or occluded processes, we manually corrected the reconstruction in ShuTu’s GUI. The reconstructions were stored in SWC format. We used ShuTu’s GUI to manually trace all visible dendritic spines. We annotated spines with thin necks and mushroom-like shapes using single SWC points projected away from the dendrite and following the spine’s neck direction. For spines with curved necks, irregular shapes or filopodia-like spines, we used multiple SWC points tracing their entire extent. We estimated spine areas in two ways. We treated spines annotated by a single SWC point as spheres attached by cylindrical necks with an average radius of 0.5 pixels (0.066 microns). For spines with multiple SWC points, we used the trapezoidal cylinders defined by every pair of consecutive SWC point. We also added a dome cap to the last SWC point. To obtain a continuous profile of spine density along each dendritic branch (Supplementary Fig. 1E), we used a rolling window of width 20 SWC points (window length fluctuated between 5 and 39 µm, with an average of 17 µm for all cells) and of step size 1 SWC point. From the continuous spine density profile, we computed the average spine density of each dendritic segment (Supplementary Fig. 1C). We excluded from the density estimation of Supplementary Fig. 1C segments branching off the soma with fewer than 10 spines and any segment shorter than 20 microns. We estimated the total dendritic area and length of each cell (Supplementary Table 1) from the reconstructions by constructing trapezoidal cylinders for each pair of consecutive SWC point. Axonal area and length were estimated in the same way, although we were only able to connect a small fraction of axon cable to the soma for 3 neurons. For 2 neurons we were not able to trace any portion of the axon (Supplementary Table 1).

We estimated the surface area of the soma of each neuron by triangulating its 3-D structure. We first used ShuTu’s GUI (Jin et al., 2019) to manually trace binary masks of the somas in all slices. Next, we traced the contours of the masks with edge detecting filters. We then took a fixed fraction of 20 equally spaced points in each contour in order to generate polygonal contours. All polygonal contours in the end had the same number of edges. Next, we applied a 3-point moving average filter along each sequence of vertex in the Z direction to smooth out rough edges. We then calculated the areas of the triangles formed by the edges and vertices of each adjacent pair of polygonal contours. We added to this number the surface areas of the first and last non-zero binary masks to represent the caps. We also corrected for the areas of the surface patches where the dendrites attach to the soma. Since these surface patches are small, we approximated them by the cross-sectional area of the first dendrite SWC point connected to the soma.

We estimated the volumes of the dendrites using 3-D binary masks which we constructed from the SWC structure. First, we re-scaled the SWC along the Z-axis to match the scale in the XY plane. Next, we interpolated the space between each connected pair of SWC points by trapezoidal cylinders with radii equal to the SWC points’ radii. The binary mask assigns ones to voxels intersecting either a SWC point or a trapezoidal cylinder and zeros otherwise. The volume of the dendrite is the fraction of active voxels in the mask multiplied by the volume of the voxel.

We estimated the volumes of the somas using an adaptation of our method for the surface areas. Starting from the polygonal contours representing the Z-stack, we used the polygons’ vertices and baricenters to split the soma’s volume into a collection of tetrahedrons. The vertices of an edge in a slice, the vertices of the closest edge in the next slice, and the baricenters of the two contours define 4 tetrahedrons for which the volume can be easily calculated. The total volume of the soma is the sum of the volumes of the individual tetrahedrons.

We approximated the volumes of the spines directly from the SWC structure. Since the spines are small, we estimated their volumes to be approximately the sum of the volumes of the individual SWC points.

### Comparative genomics of the KCNC/Kv3 (Shaw-related) potassium channel genes

To identify the full set of genes comprising this gene family in zebra finches, we first retrieved all genes annotated as KCNCs (voltage gated channel subfamily C members) from the latest RefSeq database for zebra finch (Annotation Release 106; GCF_003957565.2 assembly). We next retrieved the similarly annotated genes in other selected songbird and non-songbird avian species, observing the immediate synteny and correct cross-species BLAST alignments. To verify the correct orthology of avian genes to corresponding members of this gene family in mammals, we conducted cross-species BLAST searches, noting the top-scoring reciprocal cross-species alignments as well as conserved synteny as orthology criteria. To build cladistic trees for evolutionary inferences, we also included (as outgroups to birds and mammals) representative extant organisms from selected branches of major vertebrate groups, as appropriate, including non-avian sauropsids (crocodiles, turtles, lizards), amphibians, and bony fishes.

### Pharmacologic compounds

DL-APV, Picrotoxin, CNQX, XE991 and iberiotoxin were purchased from Tocris Biosciences (Bristol, United Kingdom). TEA and 4-AP were purchased from Sigma - Aldrich (St. Louis, Missouri, USA). α-DTX was purchased from Alomone labs (Jerusalem, Israel). AUT5 was provided as a gift from Autifony Therapeutics (Stevenage, United Kingdom).

### Statistical data analysis and curve fitting

Data were analyzed off-line using IgorPro software (Wave-metrics). Statistical analyses were performed using Prism 4.0 (GraphPad). Specific statistical tests and outcomes for each analysis performed are indicated in the respective Figure Legends and Tables. Means and SE are reported, unless otherwise noted. Electrophysiology and morphology data is described as technical replicates for individual cells while molecular data is described as biological replicates for individual birds.

## Competing Interests

None

## Data availability

A source Excel data file is included with the manuscript and a raw data file has been uploaded to the DRYAD data base (DOI https://doi.org/10.5061/dryad.1zcrjdfvs).

## Acknowledgements

We thank Autifony Therapeutics and Dr. Martin Gunthorpe for providing the AUT5 compound for this study, Drs. Manuel Covarrubias and Qiansheng Liang for thoughtful comments on experimental design using AUT5 and Dr. Cesar Ceballos for his technical recommendations regarding biocytin cell fills. Grants from NIH and NSF.

## Author Contributions

BMZ and HvG designed electrophysiology and morphology experiments. BMZ and AD acquired electrophysiology data and performed biocytin fills. AAN and CVM designed molecular biology experiments and genomic analysis. AAN acquired the molecular and genomic data. LET and DZJ designed and carried out morphometric analyses using the ShuTu software package, with help from AAN for imaging. BMZ and HvG analyzed electrophysiology data. BMZ, HvG and CVM integrated the molecular and electrophysiology data, with help from AAN and PVL. BMZ produced the first draft of the paper, HvG and CVM edited it, and all authors contributed comments and edits.

## Supplementary Information

**Supplementary Figure 1.**
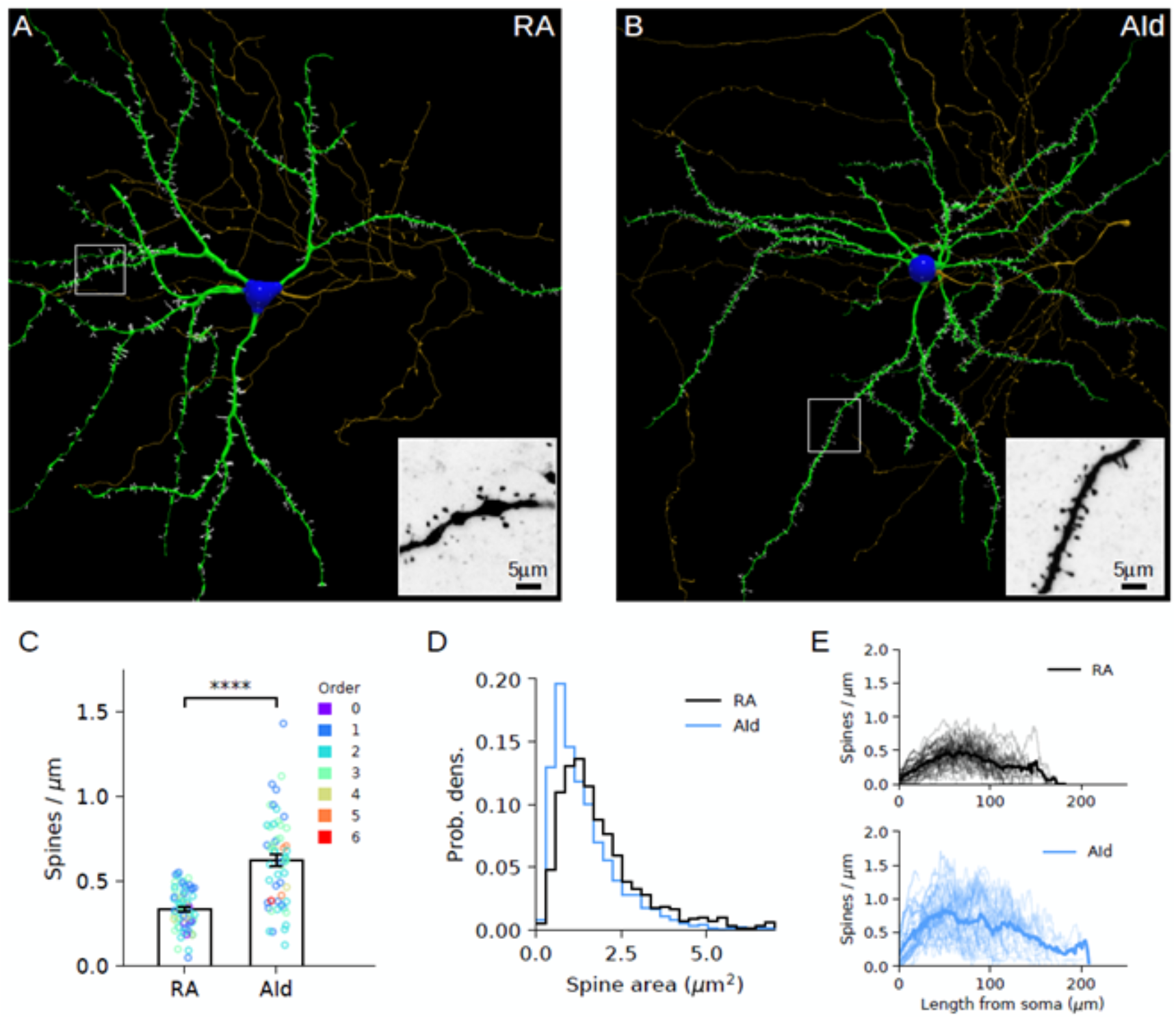
Reconstruction of neuron morphology and estimation of spine density. A & B. Top view of the morphology of a RA and an AId neuron (blue, soma; green, dendrite; gold, axon; white, spines). Insets show projections of the raw images of the regions enclosed by white boxes. Note the larger concentration of spines for the AId neuron. C. Average density of spines over dendritic segments. Circles indicate individual segments. Color indicates branching order (centrifugal ordering). Root segments with fewer than 10 spines were discarded from the analysis. Segments shorter than 20 microns were also discarded (details in Methods). Black bars indicate standard error. Welch’s t-test, t_stat_=7.75, P = 3.76 × 10^−12^, N = 62 RA and 56 AId segments. D. Probability density of individual spine areas by neuron type. Note the longer tail of the RA distribution. Kolmogorov-Smirnov two-sample test, D = 0.19 P = 7.8 × 10^−16^, N = 1194 RA and 1824 AId spines. E. Profile of spine density from soma to termini along all dendritic branches of both neuron types. N=96 RA and 95 AId branches. Averages shown in solid lines.

**Supplementary Figure 2.**
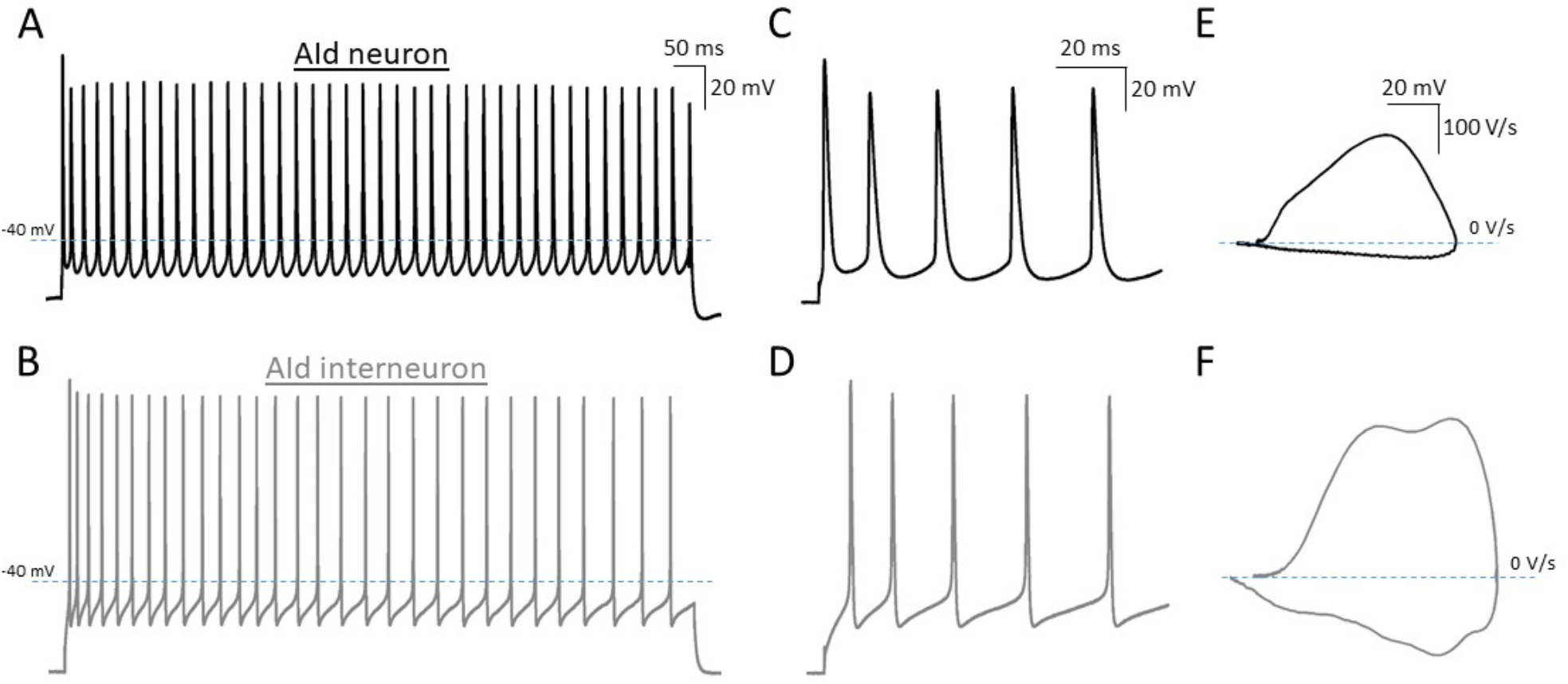
Interneurons vs. projection neurons in AId. A & B. AP trains elicited in an AId neurons (top) and a putative AId interneuron (bottom) during a 1 sec 500 pA current injection at 24°C respectively. Dashed line represents -40 mV. Scales in A. C-D. First 5 APs trains from the trains in A & B. Note the narrower APs for interneurons in the example. Scale in C. E-F. Phase plane plots from averaged APs in A & B. Scale in E. Note the larger rates of depolarization and repolarization for interneurons in the example.

**Supplementary Figure 3.**
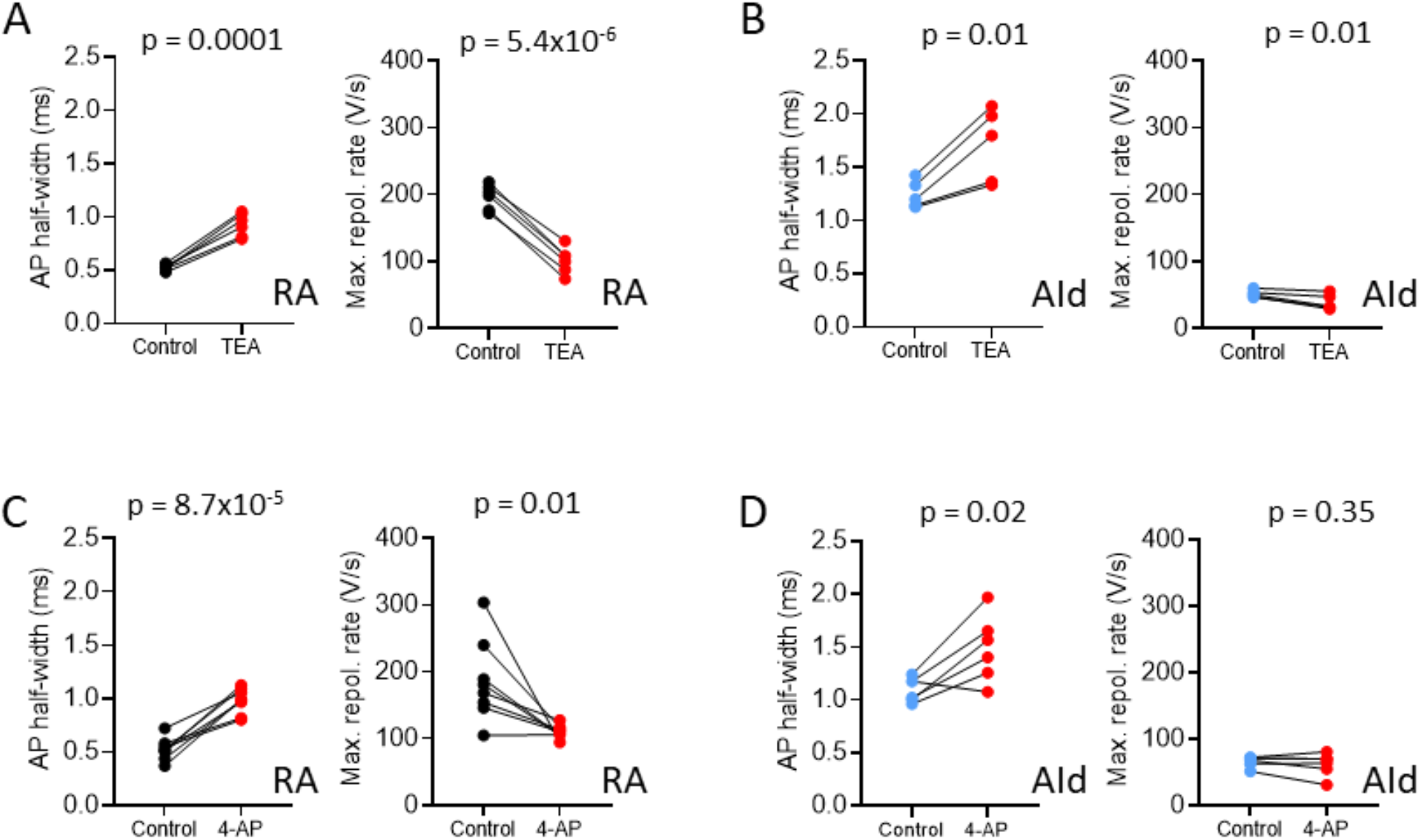
Kv3.1 antagonists broaden the RAPN and AId neuron spontaneous APs. A. Changes in AP half-width (left) and maximum repolarization rate (right) before and after exposure to 500 µM tetraethylammonium (TEA). AP half-width: Paired student’s t-test, two tailed, t_stat_ =10.90, N= 6 RAPNs; Maximum repolarization rate: Paired student’s t-test, two-tailed t_stat_= 20.28, N= 6 RAPNs. B. Same as in (A) except in AId neurons. AP half-width: Paired student’s t-test, two tailed, t_stat_= 4.51, N= 5 AId neurons; Maximum repolarization rate: Paired student’s t-test, two-tailed, t_stat_= 4.26, N= 5 AId neurons. C. Changes in AP half-width and maximum repolarization rate before and after exposure to 100 µM 4-aminopyridine (4-AP). AP half-width: Paired student’s t-test, two-tailed, t_stat_ = 8.06, N= 8 RAPNs; Maximum repolarization rate: Paired student’s t-test, two-tailed, t_stat_ = 3.28, N= 8 RAPNs. D. Same as in (C) except in AId neurons. AP half-width: Paired student’s t-test, two-tailed, t_stat_ = 3.38, N= 6 AId neurons; Maximum repolarization rate: Paired student’s t-test, t_stat_ = 1.03, N= 6 AId neurons, two-tailed. P-values are included in the graphs in A-D.

**Supplementary Figure 4.**
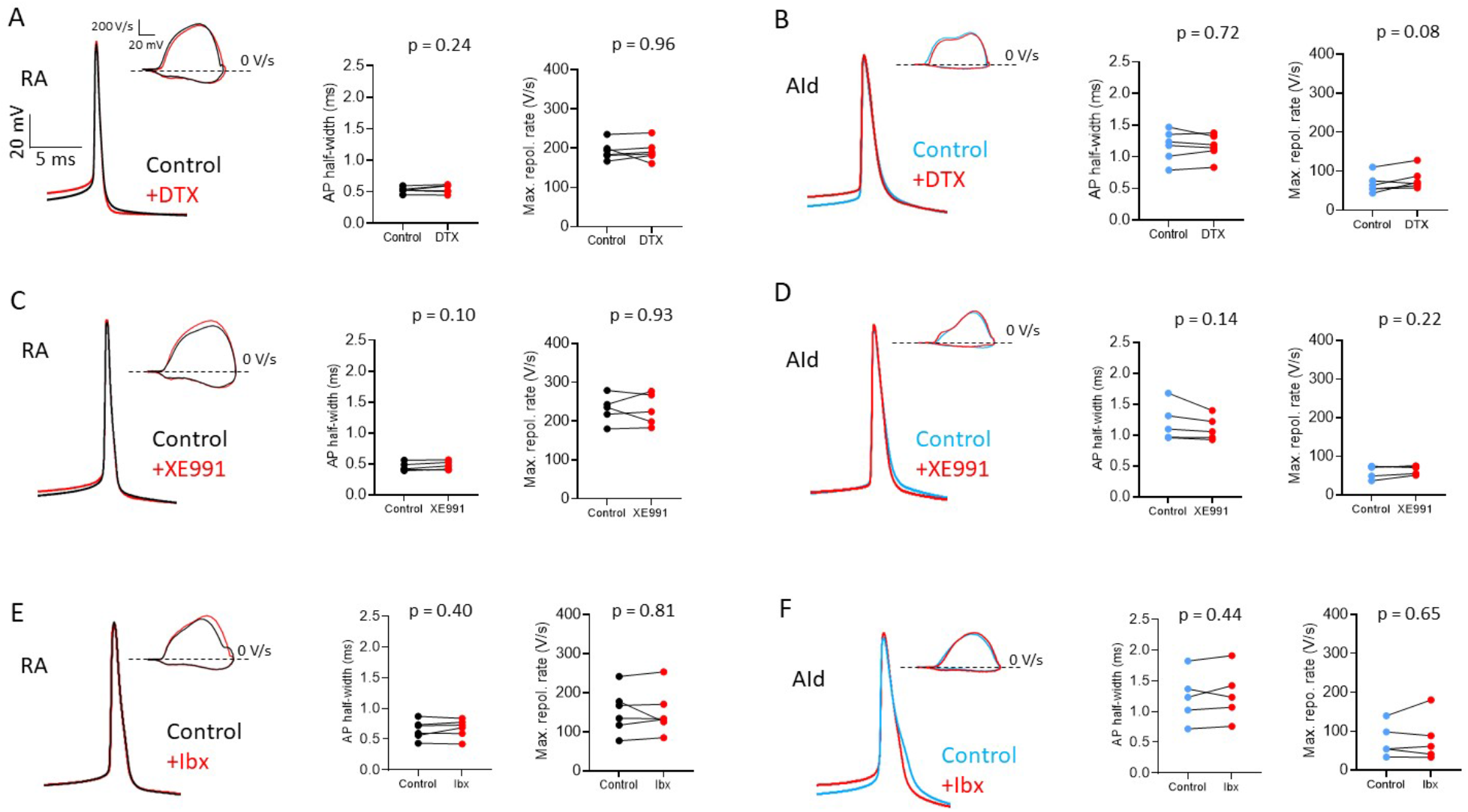
Inhibitors of TEA- hypersensitive channels do not affect RAPN and AId neuron spontaneous APs. A. Left to right: Representative averaged spontaneous AP traces, phase plane plots (inset), and changes in AP half-width and maximum repolarization rate before and after exposure to 100 nM α-dendrotoxin (DTX) for RAPNs. AP half-width: Paired student’s t-test, two-tailed, t_stat_ = 1.34, N= 6 RAPNs, two-tailed; Maximum repolarization rate: Paired student’s t-test, t_stat_ = 0.05, N= 6 RAPNs. B. Same as in (A) except in AId neurons. AP half-width: Paired student’s t-test, two-tailed, t_stat_ = 0.38, N= 6 AId neurons; Maximum repolarization rate: Paired student’s t-test, two-tailed, t_stat_ = 2.17, N= 6 AId neurons, two-tailed. C. Left to right: Representative averaged spontaneous AP traces, phase plane plots (inset), and changes in AP half-width and maximum repolarization rate before and after exposure to 30 µM XE991 for RAPNs. AP half-width: Paired student’s t-test, two-tailed, t_stat_ = 2.16, N= 5 RAPNs; Maximum repolarization rate: Paired student’s t-test, two-tailed, t_stat_ = 0.09, N= 5 RAPNs. D. Same as in (C) except in AId neurons. AP half-width: Paired student’s t-test, two-tailed, t_stat_ = 0.14, N= 5 AId neurons; Maximum repolarization rate: Paired student’s t-test, two-tailed, t_stat_ = 0.144, N= 5 AId neurons. E. Left to right: Representative averaged spontaneous AP traces, phase plane plots (inset), and changes in AP half-width and maximum repolarization rate before and after exposure to 100 nM iberiotoxin (Ibx) for RAPNs. AP half-width: Paired student’s t-test, two-tailed, t_stat_ = 0.91, N= 6 RAPNs; Maximum repolarization rate: Paired student’s t-test, two-tailed, t_stat_ = 0.26, N= 6 RAPNs. F. Same as in (E) except in AId neurons. AP half-width: Paired student’s t-test, two-tailed, t_stat_ = 0.86, N= 5 AId neurons; Maximum repolarization rate: Paired student’s t-test, two-tailed, t_stat_ = 0.49, N= 5 AId neurons. P-values are included in the graphs. Scales are the same in A-F.

**Supplementary Figure 5.**
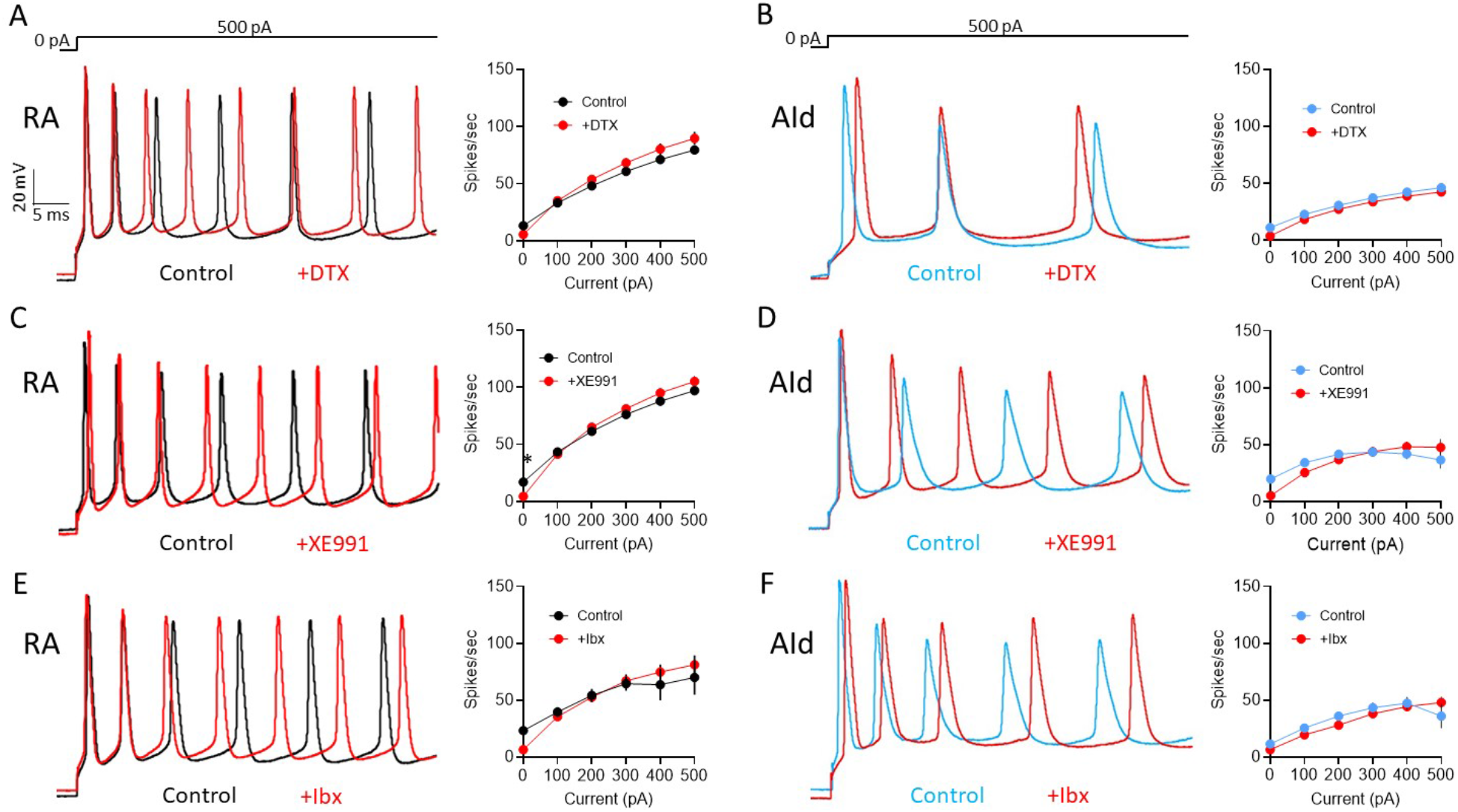
Inhibitors of TEA-hypersensitive channels do not affect RAPN and AId neuron evoked APs. A. Left to right: Representative first 50 ms of AP traces elicited by a 1 sec 500 pA current injection and average spikes/second before and after exposure to 100 nM DTX for RAPNs. Repeated Measures Two-way ANOVA; P = 1.9 × 10^−3^, F (1.269, 6.347) = 23.52, N = 6 RAPNs. B. Same as in (A) except in AId neurons. Same scale as in (A). Repeated Measures Two-way ANOVA; P = 0.3, F (1.376, 6.878) = 1.445, N = 6 AId neurons. C. Left to right: Representative first 50 ms of AP traces elicited by a 1 sec 500 pA current injection and average spikes/second before and after exposure to 30 µM XE991 for RAPNs. Same scale as in (A). Repeated Measures Two-way ANOVA; P = 6.7 × 10^−4^, F (5, 20) = 6.922, N = 5 RAPNs. * indicates P < 0.05 as determined by a Tukey’s post-hoc test. D. Same as in (C) except in AId neurons. Same scale as in (A). Repeated Measures Two-way ANOVA; P = 0.04, F (5, 20) = 2.933, N = 5 AId neurons. E. Left to right: Representative first 50 ms of AP traces elicited by a 1 sec 500 pA current injection and average spikes/second before and after exposure to 100 nM Ibx for RAPNs. Same scale as in (A). Repeated Measures Two-way ANOVA; P = 0.27, F (1.074, 5.368) = 1.546, N = 6 RAPNs. F. Same as in (E) except in AId neurons. Same scale as in (A). Two-way ANOVA; P = 0.18, F (1.686, 6.743) = 2.298, N = 5 AId neurons.

**Supplementary Figure 6.**
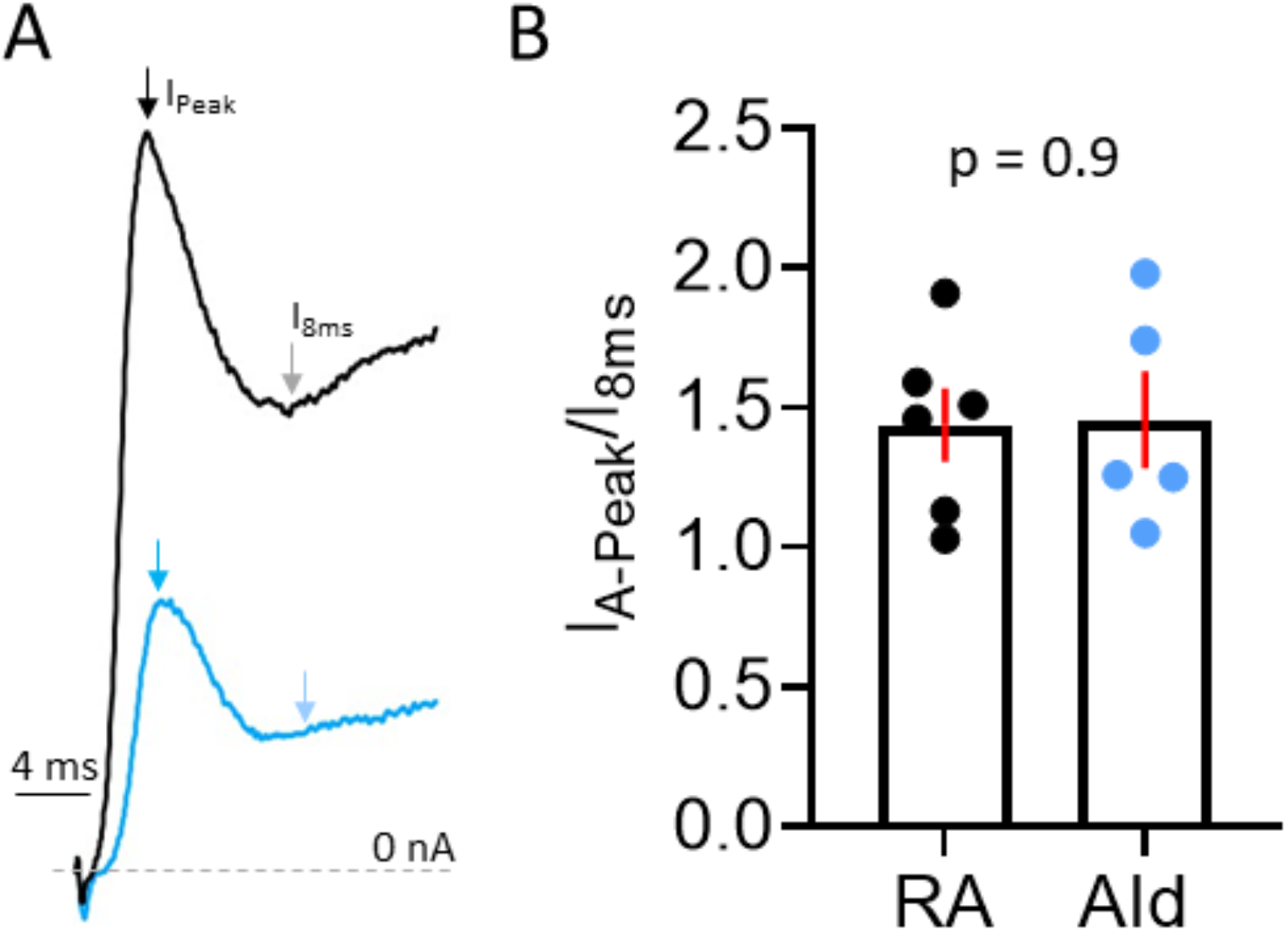
Inactivation of TEA-hypersensitive I_K+_ in RAPNs and AId neurons. A. First 20 ms of an outward TEA-subtracted I_K+_ during a step from -80 mV to 0 mV highlighting the inactivation of currents from RAPNs and AId neurons. Arrows point to the peak of the A-type component from a RAPN (black), AId neuron (blue) and 8 ms after the peak (grey and light blue arrows respectively). B. Comparison of the fold change in current from the peak of the A-type component and the current 8 ms after the peak between RAPNs and AId neurons (Student’s t-test, t_stat_ =0.08, N= 6 RAPNs and 5 AId neurons).

**Supplemental Figure 7:**
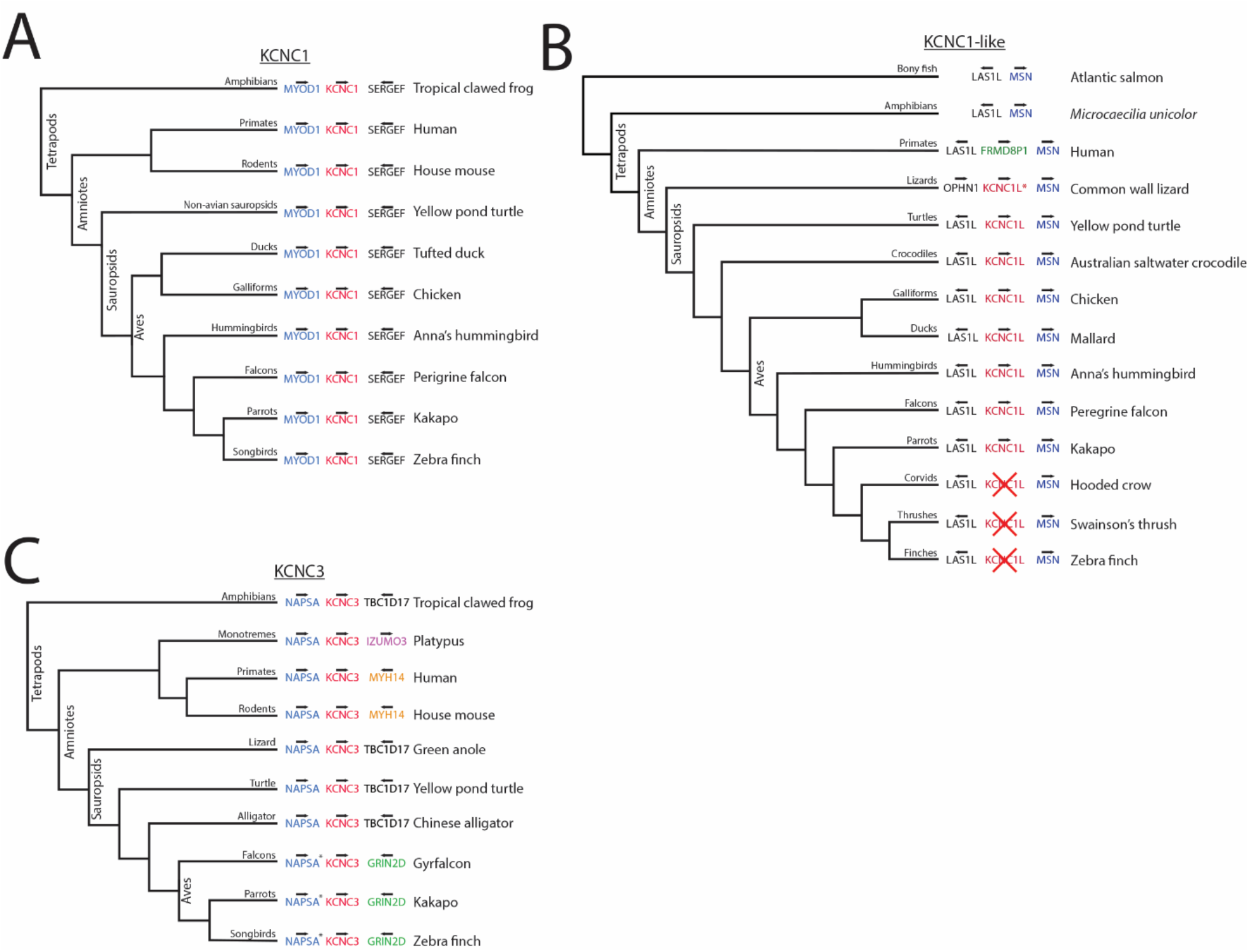
Comparative genomics of KCNC/Kv3 family members. A. Simplified cladogram for KCNC1 showing conserved synteny throughout vertebrate groups. B. Simplified cladogram for KCNC1-like paralog present in non-oscine sauropsids. Asterisk denotes a possible duplication at the KCNC1L locus in lizards. C. Simplified cladogram for KCNC3 showing the presence in birds with partial conserved synteny across vertebrate groups. Asterisk denotes NAPSA annotated as cathepsin D-like in birds. Note: For all cladograms, branch lengths are arbitrary and not calibrated for time, and only selected extant branches from major vertebrate groups are shown, to illustrate synteny conservation/divergence.

**Supplemental Figure 8:**
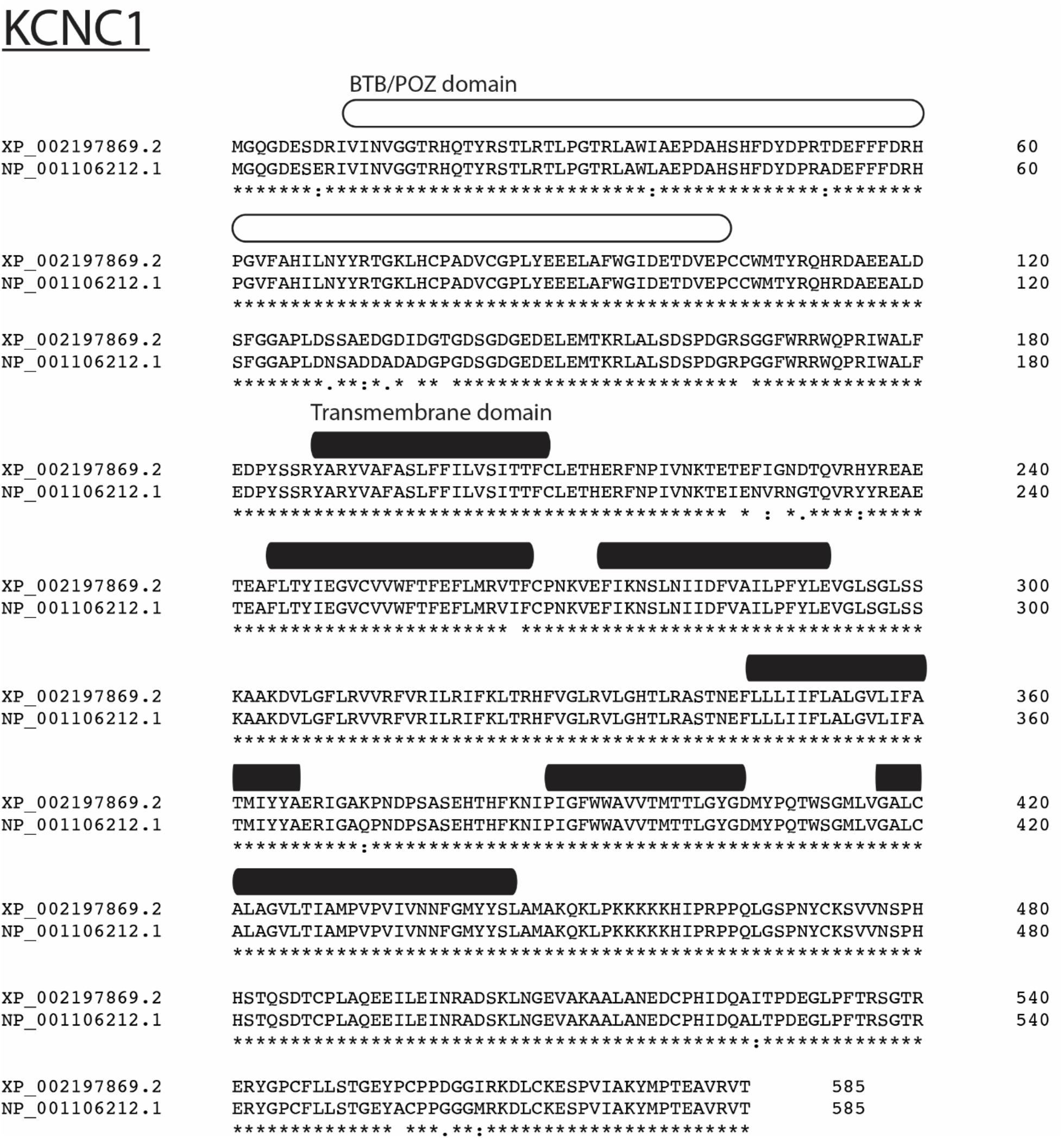
Amino acid alignment between Kv3.1 in human (top) and zebra finch (bottom). The two orthologs exhibit a high degree of amino acid conservation.

**Supplemental Figure 9:**
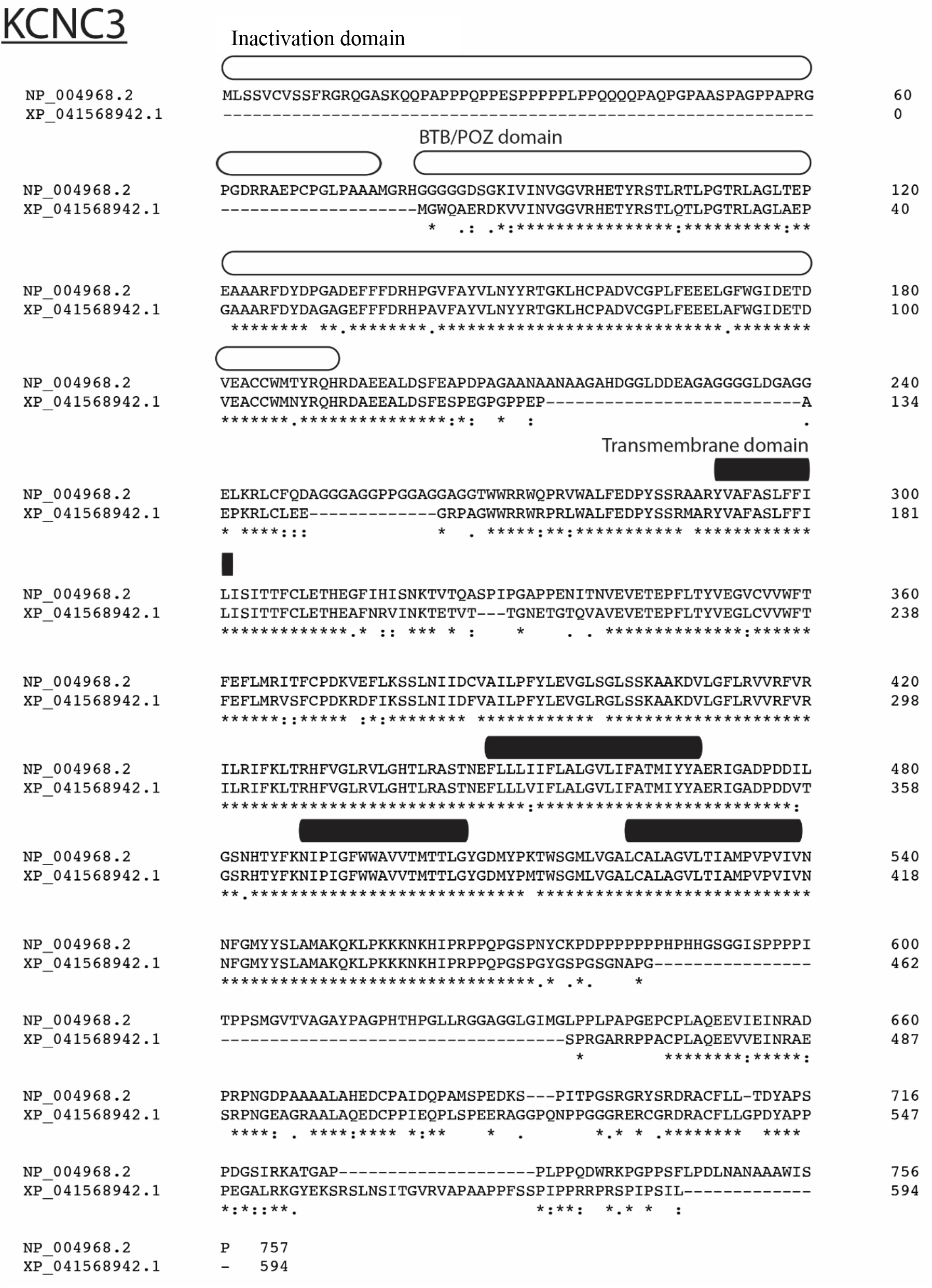
Amino acid alignment between Kv3.3 in human (top) and zebra finch (bottom). Amino acid conservation moderate and zebra finch appears to lack N-terminal domain present in humans.

**Supplementary Table 1.** Geometrical properties and spine densities measured from the reconstructions of the neuronal morphologies. Values of the axon area include only the visible portions connected to the soma. Axons excluded from volume measurements.

|  | RA1 | RA2 | RA3 | Ald1 | Ald2 |
| --- | --- | --- | --- | --- | --- |
| Soma area ($\mu\text{m}^2$) | 1125 | 599 | 915 | 721 | 733 |
| Den area ($\mu\text{m}^2$) | 5219 | 5672 | 4252 | 4143 | 6953 |
| Spine area ($\mu\text{m}^2$) | 530 | 923 | 463 | 827 | 1074 |
| Axon area ($\mu\text{m}^2$) | N/A | 99 | 286 | N/A | 1264 |
| Total area ($\mu\text{m}^2$) | 6874 | 7194 | 5630 | 5691 | 8760 |
| Total Spine # | 401 | 517 | 276 | 850 | 974 |
| Spines/ $\mu\text{m}$ | 0.25 | 0.28 | 0.2 | 0.45 | 0.39 |
| Dendritic Length | 1588 | 1883 | 1414 | 1901 | 2504 |
| Total volume ($\mu\text{m}^3$) | 4722.6 | 3155.5 | 3737.6 | 2301.2 | 3017.8 |
| Total area/volume ($1/\mu\text{m}$) | 1.456 | 2.280 | 1.506 | 2.473 | 2.903 |

## Notes

### Competing Interest Statement

The authors have declared no competing interest.

